# PYMEVisualize: an open-source tool for exploring 3D super-resolution data

**DOI:** 10.1101/2020.09.29.315671

**Authors:** Zach Marin, Michael Graff, Andrew E. S. Barentine, Christian Soeller, Kenny Kwok Hin Chung, Lukas A. Fuentes, David Baddeley

## Abstract

Localization-based super-resolution microscopy techniques such as PALM, STORM, and PAINT are increasingly critical tools for biological discovery. These methods generate lists of single fluorophore positions that capture nanoscale structural details of subcellular organisation, but to develop biological insight, we must post-process and visualize this data in a meaningful way. A large number of algorithms have been developed for localization post-processing, transforming point data into representations which approximate traditional microscopy images, and performing specific quantitative analysis directly on points. Implementations of these algorithms typically stand in isolation, necessitating complex workflows involving multiple different software packages. Here we present PYMEVisualize, an open-source tool for the interactive exploration and analysis of 3D, multicolor, single-molecule localization data. PYMEVisualize brings together a broad range of the most commonly used post-processing, density mapping, and direct quantification tools in an easy-to-use and extensible package. This software is one component of the PYthon Microscopy Environment (python-microscopy.org), an integrated application suite for light microscopy acquisition, data storage, visualization, and analysis built on top of the scientific Python environment.

## To the Editor

Localization-based super-resolution microscopy techniques such as PALM, STORM, and PAINT are increasingly critical tools for biological discovery. These methods generate lists of single fluorophore positions that capture nanoscale structural details of subcellular organisation, but to develop biological insight, we must post-process and visualize this data in a meaningful way. A large number of algorithms have been developed for localization post-processing^1, 2^, transforming point data into representations which approximate traditional microscopy images^2–4^, and performing specific quantitative analysis directly on points^1,2,5–7^. Implementations of these algorithms typically stand in isolation, necessitating complex workflows involving multiple different software packages. Here we present PYMEVisualize, an open-source tool for the interactive exploration and analysis of 3D, multicolor, single-molecule localization data. PYMEVisualize brings together a broad range of the most commonly used post-processing, density mapping, and direct quantification tools in an easy-to-use and extensible package (Figure 1). This software is one component of the PYthon Microscopy Environment (python-microscopy.org), an integrated application suite for light microscopy acquisition, data storage, visualization, and analysis built on top of the scientific Python environment^7^.

**Figure 1.**
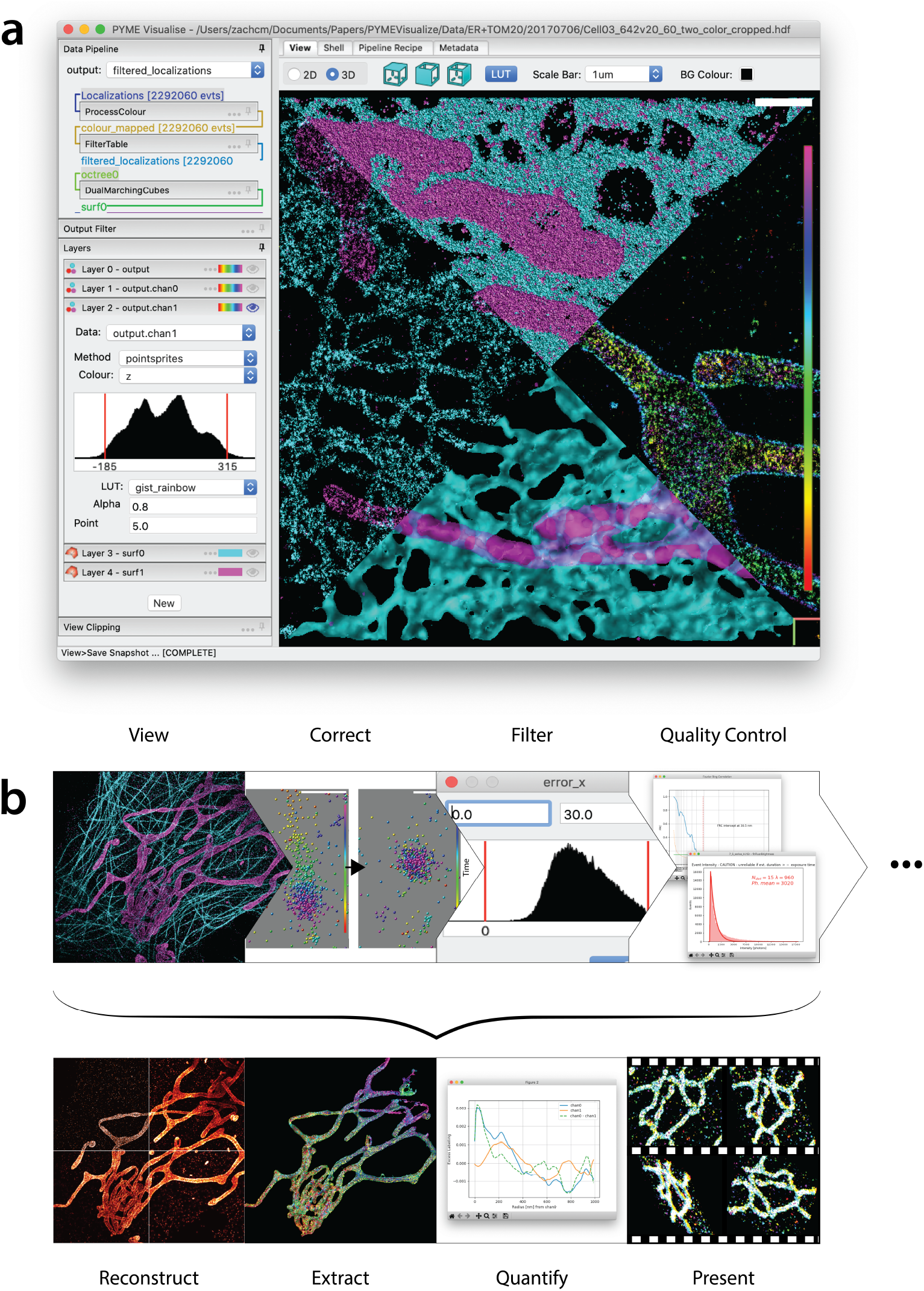
**a**. The PYMEVisualize user interface. The viewer display is divided into 4 quadrants illustrating 4 different methods of visualizing the same data set (a 2-colour dataset of the endoplasmic reticulum (cyan) and mitochondria (magenta). Clockwise from the left, the methods represent the data as the sum of Gaussians, solid spheres, a single channel (the mitochondria) colored by depth, and 3D surfaces extracted from the point-clouds. **b**. Examples of exploration, visualization, and quantification tools in PYMEVisualize. A workflow from raw events to quantification can consist of multiple steps - from an initial viewing of the localizations through corrections (fiducial-based drift correction shown), filtering on localization precision or other parameters, and the calculation of quality control metrics such as Fourier ring correlation and event photon counts. Workflows can have a wide range of endpoints such as the creation of density reconstructions (with a variety of supported algorithms), extraction of isosurfaces, quantification (analysis of pairwise distances between channels shown), and the preparation of visuals and animations. Further details about the available processing steps and the concepts underlying the PYMEVisualize implementation and user interface are described in the supplement.

PYMEVisualize is the result of over 10 years of continuous development, evolving from a simple OpenGL-based point viewer into a complete platform for the analysis of localization microscopy point-clouds, including modules for quality control, artifact correction, density image reconstruction, and quantitative analysis. It has become an indispensable tool in our laboratories and those of several collaborators, facilitating widely-varying workflows such as cluster analysis in multi-colour ratiometric dSTORM data, correlative confocal and STORM imaging, PAINT imaging of DNA origami with fiducial correction, channel registration and z-mapping for multicolor biplanar astigmatism data, extracting organelle surfaces from 4Pi-SMS images of the endoplasmic reticulum and Golgi, and generating 3D animations for use in presentations^8–11^. Its configurable processing pipeline allows for flexibility and repeatability.

We believe PYMEVisualize has the potential to become the go-to tool for the analysis and visualization of localization point data sets, akin to the status ImageJ has for pixel-based microscopy images, and that we have a duty to share it with the broader imaging community. Loading data from a wide selection of localization packages is possible through our CSV and MATLAB importers (see supplement), and a flexible plugin system makes it easy to extend. PYMEVisualize is in active development, and we welcome contributions from everyone - be it to the core code, through the creation of your own plugin(s), or simply by providing feedback about your experience. The source code is available on GitHub (https://github.com/python-microscopy/python-microscopy), and queries, feedback, or suggestions can be made directly, on the GitHub issue tracker, or using the “pyme” tag in the image.sc forum. We have included parts of the PYMEVisualize documentation - the “PYMEVisualize User Guide” and a brief tutorial (“A quick tour of PYMEVisualize”) as supplementary notes 1 and 2, respectively, and an accompanying video (supplementary video 1).

## Code Availability

Source code for PYMEVisualize is available at https://github.com/python-microscopy/python-microscopy. Compiled packages can be downloaded from https://python-microscopy.org/downloads/. For installation guidance, see the supplement.

## Data availability

The data used to generate the figures in the paper is available form the corresponding author on reasonable request.

## Acknowledgements

As a long-running project this work has received funding from a range of sources including the NIH (R01 GM118486-02, U01EB021232) the Wellcome Trust (203285/B/16/Z), the Health Research Council (NZ) and Marsden Fund (NZ). The authors wish to thank Yongdeng Zhang and Lena Schroeder for generating the ER and mitochondria and microtubule and OMP25 datasets, to thank Yongdeng Zhang, Mengyuan Sun, Ben Rollins and Joerg Bewersdorf for helpful discussions.

## Author contributions

DB created the PYME package. ZM, MG, KC, CS, AB, and DB contributed to the codebase. LF suggested features, usability improvements, and tested the software extensively. All authors contributed to the writing of this manuscript.

