## Supplementary note 1 for "PYMEVisualize: an open-source tool for exploring 3D super-resolution data"

### PYMEVisualise User Guide

**The python-microscopy team**

**Sep 28, 2020**

**7 Animation** **16**

**8 Synthetic data** **16**

**9 Editing the pipeline "recipe"** **16**

**10 Programmatic usage** **19**

**References** **22**

---

### 1 Installation

**`PYMEVisualize`** is a part of PYthon Microscopy Environment (PYME) (http://python-microscopy.org/), an open-source package providing image acquisition and data analysis functionality for a number of microscopy applications, but with a particular emphasis on single molecule localisation microscopy (PALM/STORM/PAINT etc . . . ). There are multiple routes for installation, detailed at https://www.python-microscopy.org/doc/Installation/Installation.html . The simplest installation route uses a packaged installer on Windows or macOS (see Windows instructions below).

#### 1.1 System requirements

PYMEVisualize runs on Windows, OSX, and Linux. It will run on relatively low spec machines (even a RaspberryPI), but for an enjoyable user experience we recommend:

- ~2GHz dual core CPU
- 4 GB RAM
- Hardware OpenGL support
- Wheel mouse for zooming in the interactive display

#### 1.2 Installation on Windows using executable

1. Download the latest package from http://python-microscopy.org/downloads/.
2. Double-click `python-microscopy-XX.XX.XX-Windows-x86_64.exe`.
3. If prompted with *Windows protected your PC*, click *More info* and then *Run anyway,* as shown in Fig. 1.1 a.
4. Follow instructions in the installer, leaving all options at their default.
6. When the installation is finished, locate the **`PYMEVisualise`** shortcut on the desktop. Double click it to launch PYMEVisualize, as shown in Fig. 1.1 b.

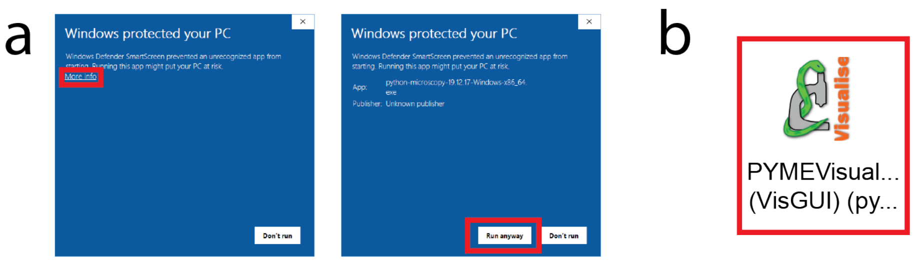

Fig. 1.1: Single-click installation and how to launch PYMEVisualize. (a)Potential Windows Defender error messages that can be safely ignored to install PYMEVisualize. (b) Double-click the **`PYMEVisualize`** logo on the desktop to start PYMEVisualize.

### 2 Data exploration

#### 2.1 Importing data

Data can be opened using the *File→Open* menu command after launching PYMEVisualize, or by specifying the filename on the command line (e.g. **`PYMEVis C:\path\to\file.h5r`**)[1] .

Three data formats are currently supported for localisation data: HDF5 (**.h5r/.hdf**), delimited text **.txt/.csv**, and matlab **.mat** files. In each case, the data should take the form of a table of values where each row corresponds to a detected single molecule event and each column corresponds to a parameter. The .txt/.csv and .mat importers are flexible and support a variety of different column layouts, with the only hard requirements being that there are columns `x` and `y` for the position of a molecule and that all columns have equal lengths. Files may contain as many other columns as they like, and columns can be in any order. To take full advantage of PYMEVisualise, the following parameters should also be included: the time/frame number at which the event was detected, the event amplitude, event width, and the estimated localisation error, accessed through the column names `t`, `A`, `sig`, and `error_x`, respectively.

##### .h5r/.hdf formats

The .h5r format is a custom format based on top of HDF5 and used by the analysis components of PYME to save localisation results and metadata. It has fixed table and column names and everything needed is read automatically from the file. It is significantly faster to read and has a smaller file size than .mat and .txt/.csv

The .hdf format is a slightly more generic HDF5 based format which has more freedom with how data is arranged within the HDF5 container. This is a good target for programs wishing to save data for use in PYME and avoid the performance issues inherent in saving as .txt.

##### Delimiter separated text (.txt/.csv)

PYME supports both tab and comma delimited text files using the .txt and .csv extensions, respectively. In both cases, the column names are defined using an import dialog (Fig. 2.1 a). It is possible to pre-populate the column names to speed up the process by adding a python style comment (signified by a leading `#`) to the first line of the file containing a list of delimiter separated column names. The dialog will still be shown for confirmation, but the correct column names should already be entered.

##### Matlab .mat files

We support MATLAB files in two formats: each column stored as a separate variable in the .mat file, or all columns in a single variable (2D array). The first format is preferred. If the variable names in the .mat file correspond to the standard variable names (`x`, `y`, `z`, etc . . . ) described in *Importing data*, .mat files will open automatically. Alternatively, an import dialog (Fig. 2.1 a) will allow mapping of column names upon import, as described in *Delimiter separated text (.txt/.csv)*. If the .mat file contains a single array, the import dialog (Fig. 2.1 b) is a little more primitive, but the same principle applies: each column needs to be given a name, and the parameters `x` and `y` must be defined. The names are specified by typing a comma separated list of parameters into the supplied box. Each of the parameters must be enclosed in double quotes, and there must be exactly the same number of parameters as there are columns in the 2D MATLAB array.

[1] You can also associate PYMEVisualise with a particular file type by using the "Open With" command in the windows explorer and locating the PYMEVis.exe (under `Scripts` in the directory you installed PYME to).

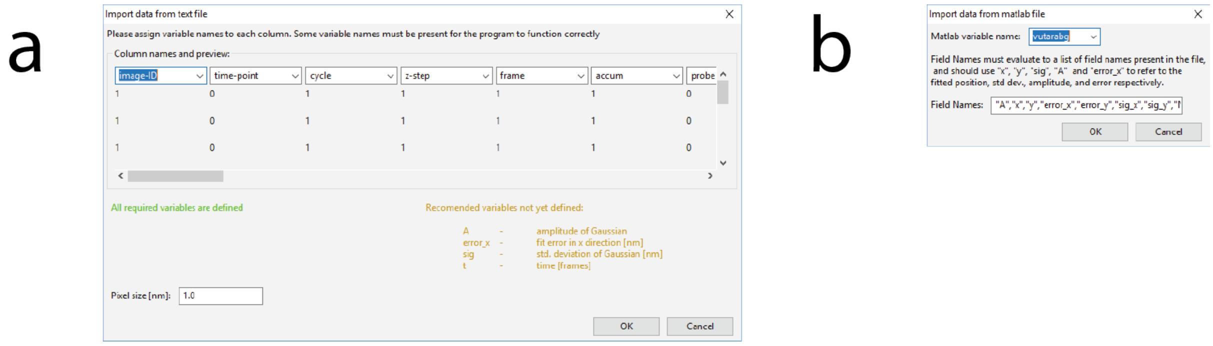

Fig. 2.1: Import dialog boxes for .txt/.csv and .mat file. (a) The dialog box that pops up when opening a text (.txt) or comma-separated value (.csv) file or a multi-column MATLAB (.mat) file. It lists a guess for each parameter name and the first ten values in that column. Columns can be renamed to match the recommended parameters not yet defined (yellow). The green text on the left indicates that required parameters (`x` and `y`) have been defined. (b) The dialog box that pops up when opening a single-array MATLAB (.mat) file. The name of the 2D MATLAB array containing localisation data is specified in the *Matlab variable name* box, and parameter names for each column within that array are specified by typing a comma-separated list of parameters into the *Field Names* box.

#### Metadata

Acquisition metadata describing camera properties, localization routines, etc., can be important for quality control and analysis. Metadata is automatically loaded from .hdf/.h5r files, and improved metadata handling for other file formats is on our TODO list. In the meantime, missing metadata can be supplied by the user in the *Shell* tab of PYMEVisualize (see *Shell*). For example, estimation of dye *photophysics* requires the `Camera.CycleTime` metadata entry (see *Photophysics*). To set `Camera.CycleTime` to 100 milliseconds, enter `pipeline.mdh['Camera.CycleTime'] = 0.100` into the shell. For more information on metadata, see http://python-microscopy.org/doc/metadata.html.

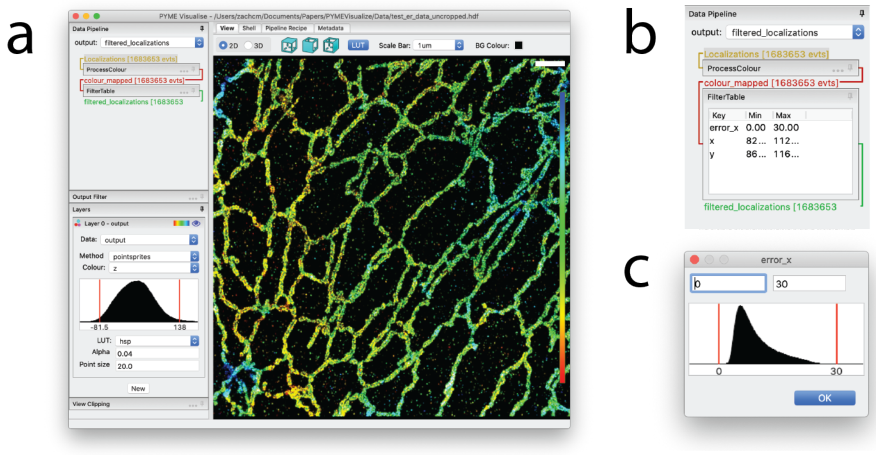

Fig. 2.2: The PYMEVisualize GUI with a loaded data set. (a) Interactive display of ~1.7 million data points from a super-resolution image of the endoplasmic reticulum in a U2OS cell, courtesy of Yongdeng Zhang and Lena Schroeder. (b) The expanded filter for this image. (c) An example editing dialog for the `error_x` filter.

Having successfully loaded a dataset, the window should resemble Fig. 2.2 a. If nothing is displayed, don't panic: the most common reason is that the filter (see *The filter* section below) is throwing away all the data points.

#### 2.2 The data pipeline

Data loaded into PYMEVisualise is processed using a configurable pipeline, accessed in the PYMEVisualize graphical user interface under the *Data Pipeline* tab (see Fig. 2.2 b for an example). By default, the pipeline loads with two sections, `ProcessColour` which extract and process colour information, if present in the input, and `FilterTable` which filters on localization precision etc ... Expanding portions of the pipeline, such as *FilterTable* (see Fig. 2.2 b, and the section below), allows for direct manipulation of their settings. Many of the additional manipulations accessible from the menus, such as drift correction and repeated localization chaining, will add steps to this pipeline. The parameters of these steps are then adjustable and will update the output in real-time. The entire pipeline can also be saved and re-loaded at a later date.

##### The filter

The filter (Fig. 2.2 b) restricts analysis and visualization to a subset of the data source. It allows specification of a valid range for each parameter, and points with parameters in these ranges are kept. The filter is used to discard erroneous events where, for example, the localization fit yielding the point picked up a noise spike or did not converge.

The filter is controlled from within the data pipeline in the sidebar, and can be expanded by clicking on *FilterTable*. Right clicking in the table gives you the option to add and, if a parameter is selected, edit or delete a parameter. Double-clicking on a parameter also enables editing. Editing parameters brings up a dialog, as shown for the `error_x` parameter in Fig. 2.2 c. A histogram of the selected parameter is displayed and the current bounds are indicated by red vertical lines. These lines can be dragged with the mouse to change the filter bounds. The filter editor (and all other histogram editors) also understand the following keys if they have focus (i.e. if the user clicks on the histogram).

| | |
|---|---|
| `m` | sets the bounds to the minimum and maximum values of the variable |
| `p` | sets the bounds to the 1st and 99th percentiles |
| `l` | toggles log scaling on y-axis |

The text editing boxes above the histogram can also be used to update parameter bounds. The filter will typically come with default bounds for `A` (the point amplitude), `sig` (the standard deviation of the fitted Gaussian), and `error_x` (the estimated error in the x position). The default values in PYMEVisualize are for imaging at ~647 nm excitation with a 1.47NA objective, and quite likely need changing. Notably, `A` will need to be changed for different intensity calibrations, and `sig` will need to be changed when working at different wavelengths.

#### 2.3 Colour channels

PYMEVisualise uses a probabilistic mechanism of channel assignment through which each fluorophore is given a probability of belonging to each of the colour channels present in the sample. Initially designed to support ratiometric imaging where colour assignments are not absolute, it is a flexible model which can also support simpler scenarios where channels are well separated or imaged sequentially. Colour assignment is performed by the `ProcessColour` pipeline module and three different methods of assigning colour probabilities are available: Bayesian channel assignment for ratiometric localisation data, temporal assignment for sequentially localised fluorophores, and pre-assignment using a `probe` column for imported data where channel assignment has already been performed. The method of colour assignment will be chosen automatically based on the file metadata and the presence of columns named either `probe` (pre-assigned), or `gFrac`[2] (ratiometric). Under the hood, these all feed into the probabilistic colour model resulting in special `p_<channel_name>` columns. If multiple color channels are detected, PYMEVisualize will automatically generate layers (see *Interactive display*) for each color channel when the file is loaded, in addition to the standard layer showing all points. See *Ratiometric colour settings* and *Isolating a single channel for processing* for details on ratiometric colour processing and channel extraction for non-colour aware processing routines.

[2] Corresponding to the ratio of short channel to total intensity for a single event.

#### 2.4 ROI selection / the "Output Filter"

The "Output Filter"[3] is located immediately below the data pipeline. It is similar to the filter within the pipeline, but operates after all other processing steps and immediately before display. Its primary use is for cropping the data to a smaller spatial ROI by adding filters on the `x` and `y` parameters. Rather than manually creating and setting these filters, a selection can be made by clicking and dragging with the left mouse button within the view tab (a yellow selection rectangle should be shown), and then clicking on *Clip to Selection* in the *Output Filter* pane (or pressing `F8`). The ROI can then be cleared by clicking the same button (or by pressing `F8` again).

#### 2.5 Interactive display

The processing pipeline feeds into the interactive display (Fig. 2.2 a). By default the display shows a single **"Points"** layer which renders the processed localisations as a point cloud. Points layers (see, e.g. Fig. 2.2 a) support a number of different display modes, from simple dots, through shaded spheres, to transparent Gaussians (point sprites), which provide a real-time approximation to the popular Gaussian reconstruction mode. Points can be coloured by any of the fitted parameters (via the *Colour* dropdown), with a variety of different look up tables (*LUT*) and with adjustable size and transparency. Extra layers can be added to simultaneously visualise different steps in the processing pipeline, colour channels, or data types. In addition to the **Points** data type, there are layers for rendering triangular meshes/surfaces, octrees, single particle tracks and voxel-based image data.

The display can be zoomed in and out using the mouse wheel, and panned by dragging with the right mouse button. Choosing *View→Fit* from the menu will reset the display such that the whole data set fits within the display window. Pressing `C` recenters the data bounding box on the current view. A scale bar and color lookup table are on the right of the display window.

### 3 Data correction and quality control

#### 3.1 Chaining

A fluorophore that is on for multiple frames in the raw data will appear as a series of localizations at sequential times. To group localizations close in space and time into single events, run *Corrections→Chaining→Find consecutive appearances*. A dialog will appear allowing chaining options to be set.

**`class FindClumps`**

> **Clump radius** is the maximum spatial distance between chained localizations. The default is twice the localization's lateral fit error (a $2\sigma$ should correctly link 95% of localisations). '
>
> **Time window** is the maximum temporal distance (in frames) allowed between chained localizations.

Pressing *OK* in the dialog will then identify which localizations are likely members of a chain, but will not replace the members of the chain with a single grouped/averaged localisation. This is done in a separate step, *Corrections→Chaining→Clump consecutive appearances*.

[3] This name is historical, and refers to a time when this was the only filter in the workflow. It will probably be renamed to ROI at some point in the future.

#### 3.2 Drift correction

PYMEVisualise supports 3 forms of drift correction out of the box, with additional algorithms available as plugins. The builtin methods are as follows:

**Fiducial based drift correction** This uses fiducials localised along with the blinking molecules to correct drift, and assumes that the fiducial localisations are present in a different dataset to the single molecule localisations (as optimal detection, background subtraction, and fitting settings are likely to be different for fiducials and molecules). If the data was analysed using PYME, both these datasets should be in the same file, and running fiducial based correction should be as simple as selecting *Extras→Fiducials→Correct* from the menu (and potentially entering the fiducial diameter to permit filters to be set accordingly). If fiducial and single molecule datasets are not in the same file, you will need to load the localisations first and then run *Extras→Fiducials→Load fiducial fits from 2nd file* to load the fiducial fits. The algorithm extracts fiducial traces, and aligns and averages the traces from multiple fiducials (weighted by localisation precision). It tolerates small gaps in the fiducial traces as long as not all fiducials traces are broken in the same frame. After correction is complete, *Extras→Fiducials→Display Correction Residuals*, will show the residuals (error between each fiducial and the average correction) which gives an indication of correction quality.

**Autocorrelation based drift correction** Accessed as *Corrections→Autocorrelation based drift correction*, this is essentially an implementation of the algorithm described in the supplement of [huang2008], dividing the localizations into overlapping time blocks, and performing autocorrelation between those blocks.

**Transmitted light correction** This relies on drift measurements made during image acquisition using a transmitted light channel (see [mcgorty2013] - our implementation does not assume, however, that correction of x-y drift is necessarily performed in real-time, rather saving the recorded drift values along with the image data), and requires suitable drift *event* data in the input files.

#### 3.3 Fourier Ring Correlation

Fourier ring correlation (FRC) is an established technique for estimating the resolution of localization-based images [nieuwenhuizen2013]. To use FRC to estimate resolution in PYMEVisualize, first select *Extras→Split by time blocks for FRC*. The will create 2 fake colour channels, `block0` and `block1`, based on dividing localisations in time with a temporal block size set using the dialog in Fig. 3.1 a (for multicolour data it will split the existing color channels in 2 so you will get double the number of colour channels, e.g. `chan0_block0`, `chan0_block1`, `chan1_block0`, `chan1_block1`).

Render images by choosing *Generate→Histogram* from the menu and selecting the two blocks corresponding to the color channel of interest (the FRC module currently assumes a single color), as shown in Fig. 3.1 b. Note that in principle any image generation method (see *Image Reconstruction*) can be used, but histogram rendering is probably the best for a pure resolution assessment.

In the rendered image window, choose *Processing→FRC* and select the renderings of the two time blocks to compare as in Fig. 3.1 c. An FRC plot like Fig. 3.1 d will appear, quantifying resolution.

#### 3.4 Photophysics

For a given image, it is possible to estimate the photophysics of the dye or fluorescent protein used in acquisition. To do this, first run through the clump detection part of the chaining procedure described in *Chaining*. Then select *Analysis→Photophysics→Estimate decay lifetimes*. This will display three graphs, shown in Fig. 3.2, indicating the fluorescence decay rate of the fluorophore, the mean number of fluorophores in an ON state per second throughout the duration of imaging, and the mean number of photons per frame.

Note that the metadata setting `Camera.CycleTime`, which is the integration time of the camera used to collect the raw localization data, must be present in order to analyze photophysics. See *Metadata* for details on how to ensure this metadata is present.

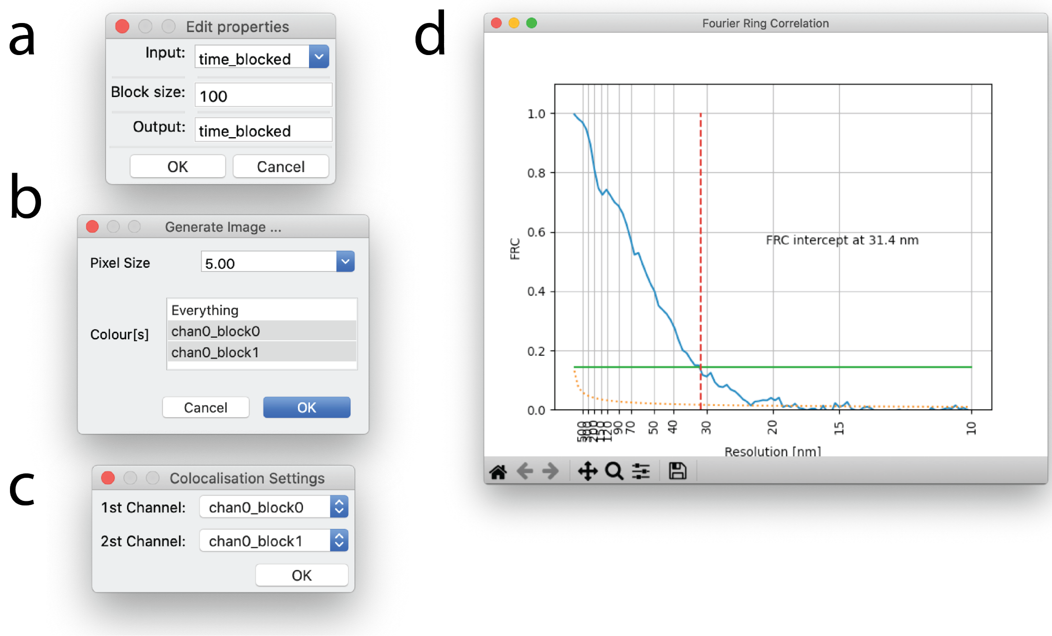

Fig. 3.1: Dialogs and plots in the Fourier ring correlation pipeline. (a) Dialog for *Extras→Split by time blocks for FRC*, used to set FRC time block size. (b) Histogram generation dialog window. Pixel size is set to 5 nm and FRC `block0` and `block1` are selected for rendering. (c) Dialog for *Processing→FRC*, indicating blocks to compare for FRC. (d) FRC plot for image shown in Fig. 2.2 a.

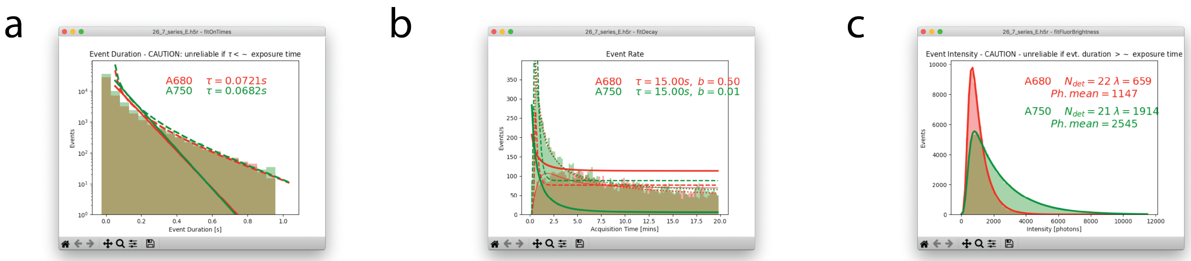

Fig. 3.2: Plots generated from running *Analysis>Photophysics>Estimate decay lifetimes* on data shown in *Ratiometric colour settings*. (a) Estimation of fluorophore decay rate, indicated as $\tau$ in the upper right of the plot. (b) Estimation of mean number of fluorophores in an ON state per second throughout the duration of imaging, indicated as $\tau$ in the upper right of the plot. (c) Estimation of the mean mean number of photons per fluorophore in the ON state, indicated as *Ph. mean* in the upper right of the plot.

### 4 Image Reconstruction

In many cases it is desirable to reconstruct a density image analogous to a more conventional voxel based dataset such as would be acquired by confocal microscopy. PYMEVisualise supports a number of different image reconstruction algorithms, which can be found under the *Generate* menu. The following methods are supported.

**Histogram** A histogram of localisation positions with a specified bin size. The simplest possible reconstruction technique.

**Gaussian** The popular reconstruction method which creates a density image by summing Gaussians at each localisation. By default, the estimated localisation error is used to determine the width of the Gaussians (as introduced in [betzig2006]), but any of the fitted parameters can be used. Using the fitted event width (`sig`) instead is a simple way of generating synthetic diffraction limited images.

**Jittered triangulation** Described in [baddeley2010], the jittered triangulation method performs a local density estimate based on a Delaunay-triangulation of the localisation data. When compared to Gaussian rendering it gives less weight to stochastic features resulting from only a few localisations and generally gives better quality segmentations when thresholded in subsequent processing. In the limit of high emitter density, it also gives better resolution (although practical emitter densities are seldom high enough for this to be relevant).

The variable which dictates the jitter magnitude can be selected, and defaults to a measure of the distance between a point and its neighbours. The number of samples to average defaults to 10.

In addition to jittering, it is also possible to smooth the triangulation by averaging several triangulations performed on Monte-Carlo subsets of the point positions. To try this out, set the multiplier for the jitter to 0 and set the MC subsampling probability to less than 1 (~ 0.2 is probably a good start).

**Quadtree** Also described in [baddeley2010], the quadtree method renders a quadtree where the intensity of each leaf of the tree is proportional to the density of points within that leaf, dividing bins when the number of localisations contained is greater than the *Max leaf size* parameter. This leads to adaptive bin sizing where areas of the image that are localisation poor have large bins and localisation dense regions have small bins. A convenient way to think of the quadtree method is that it yields an approximately constant signal to noise (of $\sim \sqrt{\frac{max\_leaf\_size}{2}}$) across the image regardless of local point density. Like the jittered triangulation method, it helps avoid some of the visual and segmentation artifacts obtained using the Gaussian method when sampling density is low.

3D versions of the histogram, Gaussian, and Jittered Tirangulation methods are also available. These generate a volumetric stack rather than a 2D image.

After an image is generated, it will pop up in a new window (Fig. 4.1 b). Colour scaling in the viewer can be adjusted by expanding the *Display Settings* tab to the right of the image. When viewing multi-colour images, individual channels will appear in separate tabs, along with a composite tab in which the channels are overlaid. Generated images may be saved as raw values (*File→Save as*) suitable for quantitative analysis in downstream software, or exported as scaled and colour-mapped snapshots (*View→Export Current View*) for inclusion in presentations or publications. The default formats are floating point TIFF for raw data and PNG for snapshots.

---

**Note:** Some versions of ImageJ/FIJI do not load floating point TIFF (and therefore our exported images) correctly, although the Bio-Formats importer does. If an exported .tif looks weird in ImageJ, try opening with the Bio-Formats importer.

---

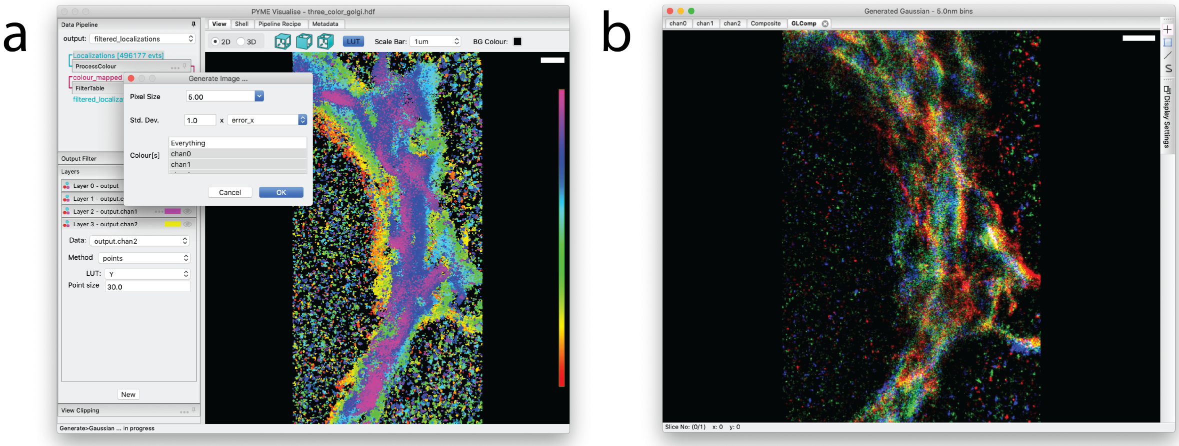

Fig. 4.1: 2D Gaussian rendering of a 3-color super-resolution image of *cis*, *medial*, and *trans*-Gogli. (a) A *Generate Image…* dialog specifying a pixel size of 5 nm, a standard deviation of `error_x` nm for each rendered Gaussian, and a request for renderings of all 3 colour channels. (b) An image viewer displaying a composite of the rendered colour channels. Individual channels are accessible via tabs (`chan0`, `chan1`, `chan2`) in the image viewer. Clicking on *Display Settings* will reveal a histogram that can be used to adjust colour channel contrast, among other display tools.

### 5 Surface extraction

#### 5.1 Isosurfaces

Isosurfaces are a common tool for visualising volumetric voxel data sets such as those produced by confocal microscopy. The algorithms and software tools used to generate isosurfaces for confocal can be applied to super-resolution images after performing a 3D density reconstruction (see *Image Reconstruction*). This indirect approach, however, has a number of disadvantages. To capture detail in the data sets generally requires the use of a small reconstruction voxel size, resulting in exceptionally large datasets. A 10x10x10 µm super-resolved volume with a 5 nm pixel size would give rise to an 8 gigavoxel (32 GB) reconstructed volume. This represents a major computational challenge, in practice limiting such reconstructions to small ROIs and often smoothed and down-sampled data. A second limitation is the need to choose this voxel size in advance. Due to the stochastic nature of localisation microscopy, choosing an appropriate reconstruction voxel size is not a trivial problem - different parts of the image could well have a different optimal voxel size.

In PYMEVisualise we have implemented algorithm which permits isosurfaces to be extracted much more efficiently from point datasets without a conventional image intermediate. Our algorithm initially places points into an octree data structure [meagher1980] (Fig. 5.1 b). We then cut / truncate the octree at a given minimum number of localisations per octree cell (equivalent to a minimum signal to noise ratio (SNR) - see [baddeley2010]). This has the effect of dividing the volume into cubic cells with a size which adapts to the local point density. Cells will be large in areas with few localisations, and small in areas which are localisation dense. The result is a volumetric data structure that contains the same information as a fully sampled reconstruction but with a lot less elements. We calculate a local density of localisations in each cell and then run the Dual Marching Cubes ([schaefer2005]) algorithm on this with a given density threshold (Fig. 5.1 c).

The algorithm for isosurface generation is accessible from the menu as *Mesh→Generate Isosurface*. This will construct the octree over which the isosurface is calculated and then display a dialog (Fig. 5.2 a) allowing parameters of the isosurface generation to be adjusted. The parameters are as follows:

**`class DualMarchingCubes`**

**Parameters**

- **`Input`** – the octree name (do not modify)
- **`NPointsMin`** – the leaf size (number of localisations) at which we truncate the octree.

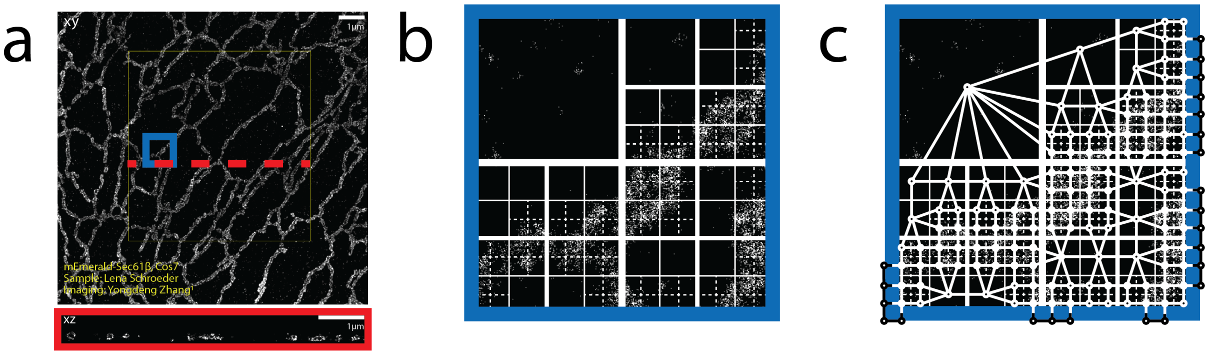

Fig. 5.1: Octree generation. (a) The 3D dataset from Fig. 2.2 requires 900 megavoxels to sample at 5 nm intervals. To reduce memory use, we sample with a sparse octree. (b) Birds-eye view of an octree used to generate an isosurface, overlaid on a subregion of the dataset shown in Fig. 2.2 a. (c) A birds-eye view of a dual grid used to generate an isosurface, overlaid on a subregion of the dataset shown in Fig. 2.2 a. Each vertex of the dual grid the center of an octree leaf.

A higher value increases the SNR at the expense of resolution.

- **`ThresholdDensity`** – the threshold on density (in localisations/nm^3) at which to construct the isosurface.
- **`Remesh`** – improves mesh quality by subdividing and merging triangles such that triangles are more regularly sized and the number of connections at each vertex is more consistent. This improves both appearance and the reliability of numerical calculations on the mesh (e.g. curvature and vertex normals). Disable when experimenting with thresholds to improve performance.
- **`Repair`** – will patch holes in the mesh (usually not needed).

#### 5.2 Spherical harmonics

Another method of surface extraction from point data sets is to fit spherical harmonics (*Analysis→Surface Fitting→Spherical Harmonic Fitting→Fit Spherical Harmonic Shell*). In contrast to the isosurface method (which simply thresholds on density) spherical harmonic fitting assumes that points lie on a surface. Because it is model based it is much better constrained and can extract accurate surfaces from significantly sparser datasets. It is ideally suited to the extraction of the cell nuclear envelope based on a lamin or NPC staining, but is also applicable to other "blobby" structures which are shell-labelled [singh2011]. When multiple objects are present in a field of view, these will need to be segmented first.

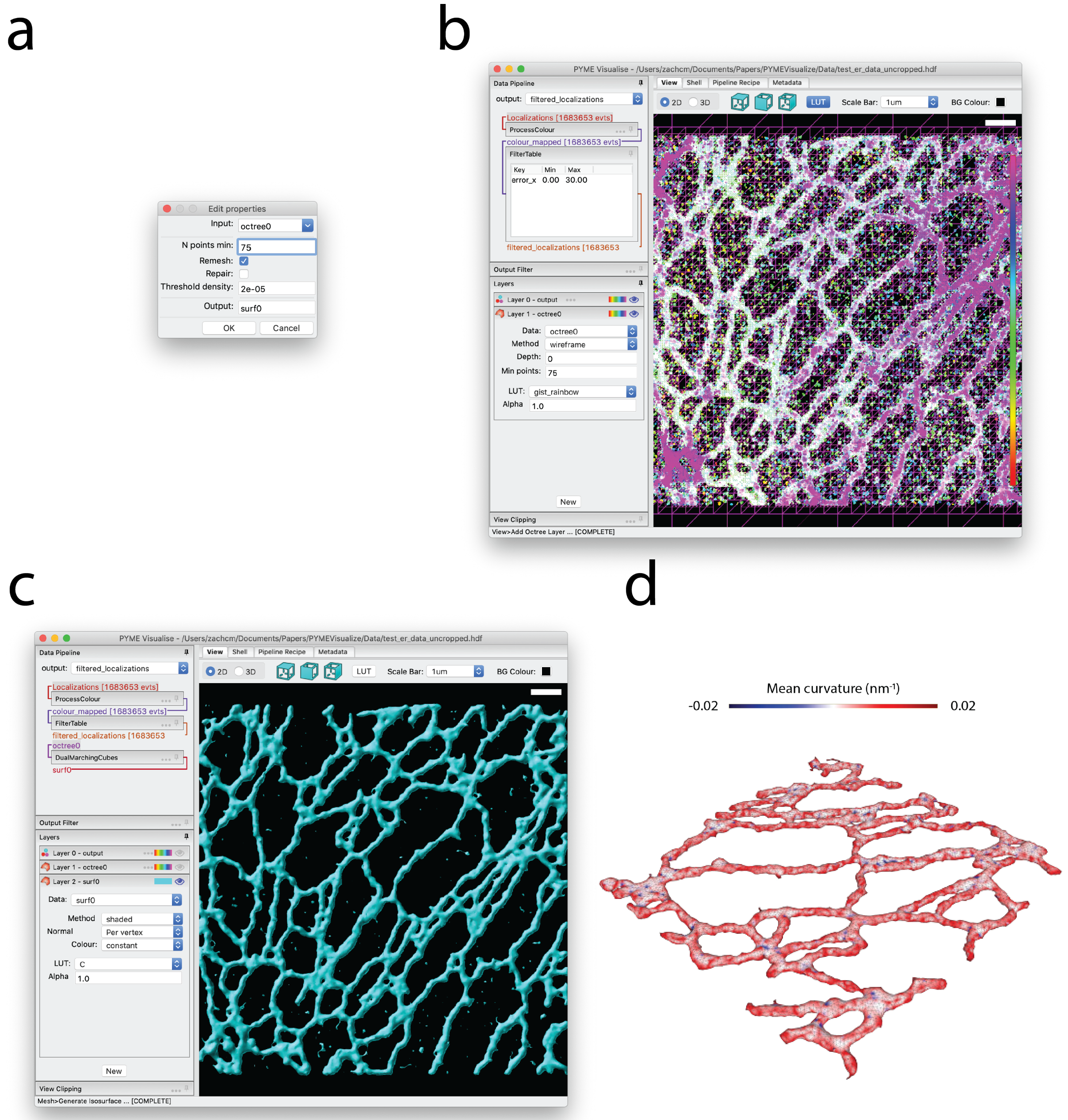

Fig. 5.2: Isosurface generation. (a) Dialog box for isosurface generation. (b) Birds-eye view of the octree layer used to generate isosurface, overlaid on the original dataset shown in Fig. 2.2 a. (c) Birds-eye view of the generated isosurface. (d) Middle portion of the isosurface in c, colored by mean curvature and rotated for perspective.

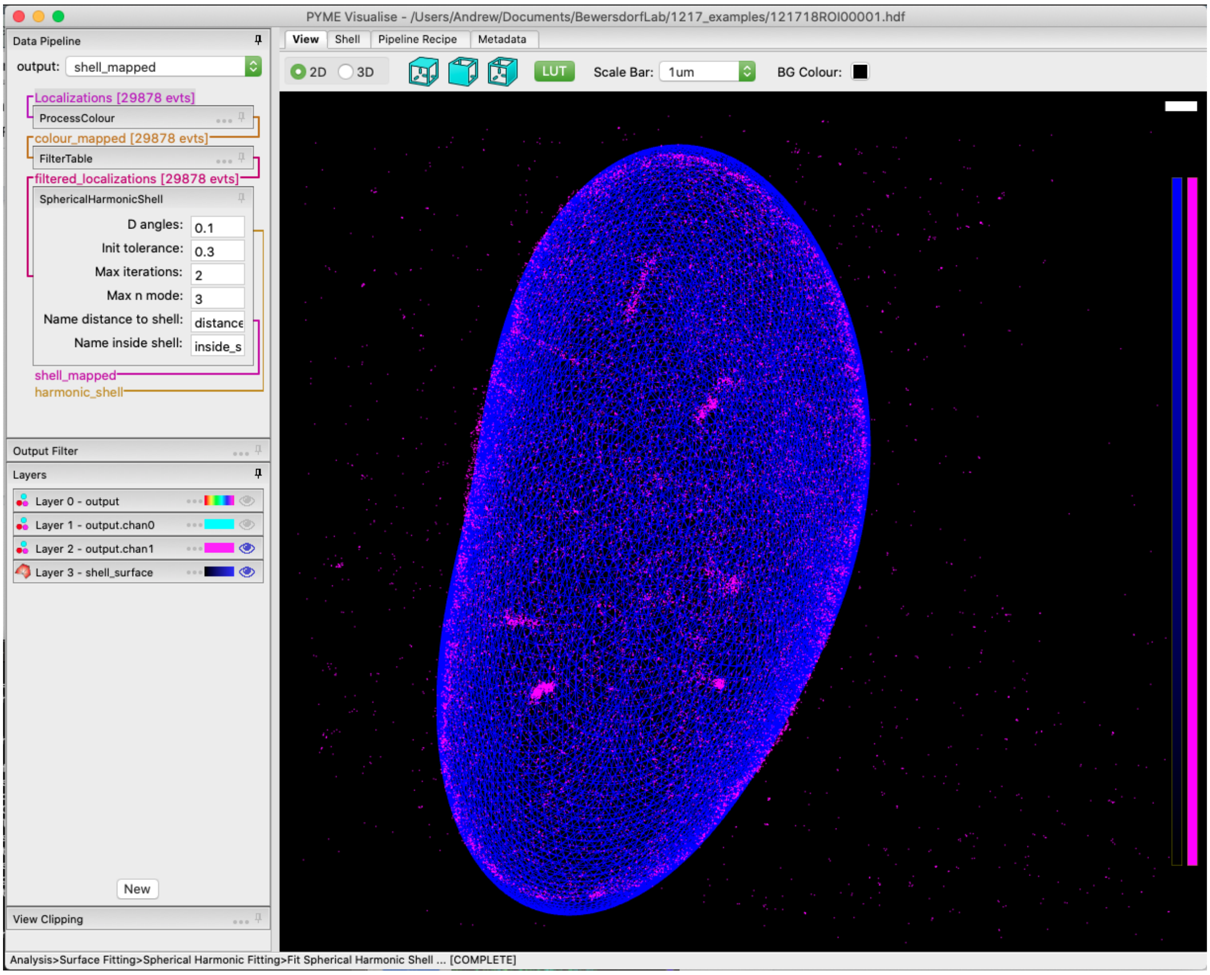

Fig. 5.3: Data from [barentine2019].

#### 5.3 Mesh manipulation

A number of operations are possible on meshes generated using either isosurfaces or spherical harmonics. These meshes can be colored by variables, such as `x`, `y`, `z`, and curvature, as in Fig. 5.2 d. Meshes can be exported to STL or PLY format, suitable for importing in other software or 3D-printing. There are also a growing number of analysis options (e.g. *Mesh→Analysis→Distance to mesh* which calculates the signed distance between localisations and the mesh) which operate on the meshes.

### 6 Quantification

PYMEVisualise includes several quantitative analysis routines, operating both directly on localisation data and on reconstructed images. An incomplete description of some of the most well used options is given below.

#### 6.1 Pairwise distances

Pairwise distance histograms are a powerful metric calculated directly from the point data set without the need for reconstruction. They measure the distribution of distances from each point to every other point and can be used for cluster analysis, either through the raw pairwise distance histogram (*Analysis→Clustering→Pairwise Distance Histogram*) or using derived measures such as Ripleys K & L functions (*Analysis→Clustering→Ripley's K/L*) [kiskowski2009], [nicovich2017]. When applied between two colour channels, they provide a co-localisation (or co-clustering) measure (*Analysis→Pointwise colocalisation*) [coltharp2014]. It is also possible to infer some shape features from the pairwise distance histogram [osada2002], [curd2020].

To the best of our knowledge, the algorithms used for calculating pairwise distances histograms in PYMEVisualise are significantly more efficient than existing published options. This is achieved by calculating the histogram on the fly and never storing the full pairwise distance matrix. To give a rough idea, we are 20x faster than the already optimised `numpy.histogram(scipy.spatial.pdist(...))` (or equivalent MATLAB calls). We also use several orders of magnitude less memory (memory demand scales as $O(N)$ rather than $O(N^2)$ as for `pdist()`) meaning that we can feasibly compute pairwise distance histograms from much larger datasets (100k points requires ~1.6MB as opposed to ~1GB).

For completeness, it is also possible to calculate a histogram of nearest neighbour distances (*Analysis→Clustering→Nearest Neighbor Distance Histogram*). Whilst somewhat easier to interpret, these tend to be much more susceptible to noise than the full pairwise distance histogram so it is preferable to use the latter where possible.

#### 6.2 Single-molecule tracking

PYMEVisualise offers a couple of particle tracking algorithms. The simplest option, suitable for well separated particles, is accessed as follows. Open a dataset containing at least the variables `x`, `y`, and `t` (see *Importing data*). Navigate in the menu to *Analysis→Tracking→Track single molecule trajectories*. A dialog box will pop up as shown in Fig. 6.2 a. The maximum distance between two consecutive points which will still be connected as a track is given by `ClumpRadiusScale*ClumpRadiusVariable` where `ClumpRadiusVariable` can be any of the fitted parameters/columns or a constant. For particle tracking it makes sense to leave this at it's default of a constant 1 nm and just alter `ClumpRadiusScale`. `ClumpRadiusScale` should be set to a value that is greater than the maximum distance a particle is expected to move within one frame[4], but less than the distance between separate particles. `Min clump size` refers to the minimum number of points that need to be in a track. `Time window` is the maximum distance in time between any two consecutive points in a track. Once tracked, the trajectories will display in a tracks layer (see *Interactive display*) as in Fig. 6.2 b. Tracks can be optionally colored and/or filtered by variables from the original dataset, by track unique identifier, by track length, and by instantaneous velocity. A slightly more sophisticated tracking algorithm, suitable for denser tracking scenarios, which can use the z-position along with additional features such as particle brightness and shape to improve linkages is available by manually adding the `TrackFeatures` module to the pipeline (see *Editing the pipeline "recipe"*).

[4] Or twice the localisation precision if the molecules are super slow moving and their expected motion is less than this.

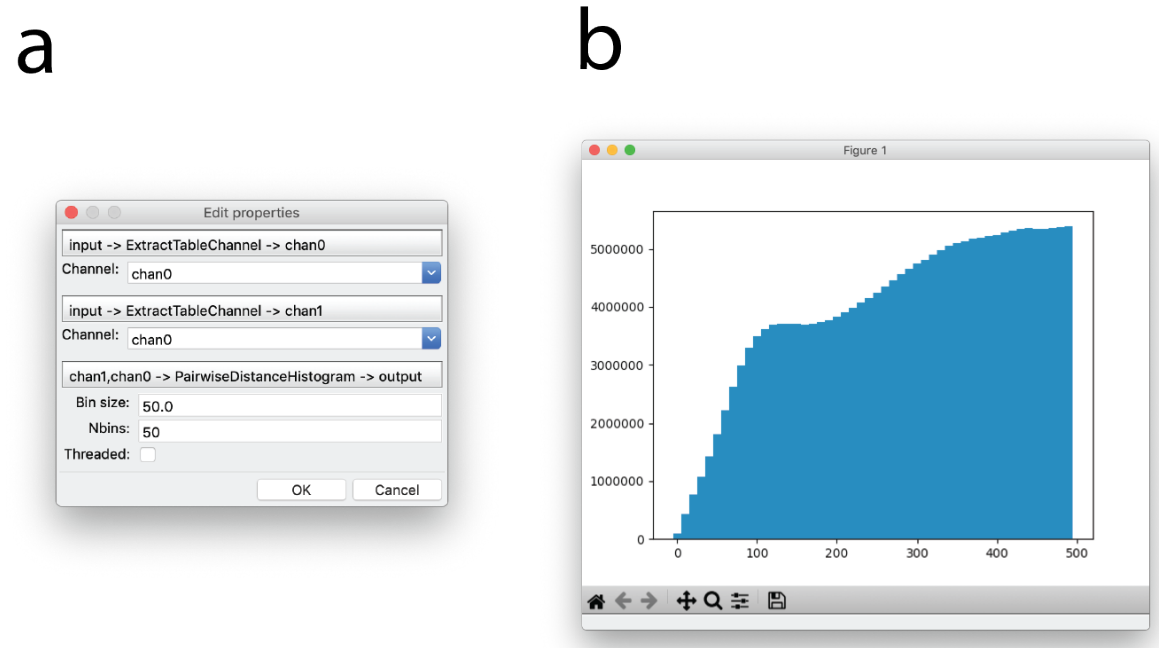

Fig. 6.1: Pairwise distance histograms. (a) Dialog for generating pairwise distance histograms. (b) Example pairwise distance histogram of sub-ROI of the image in Fig. 2.2 a with a bin size of 10 and 50 bins. Distance in nanometers is plotted on the x-axis and counts are plotted on the y-axis.

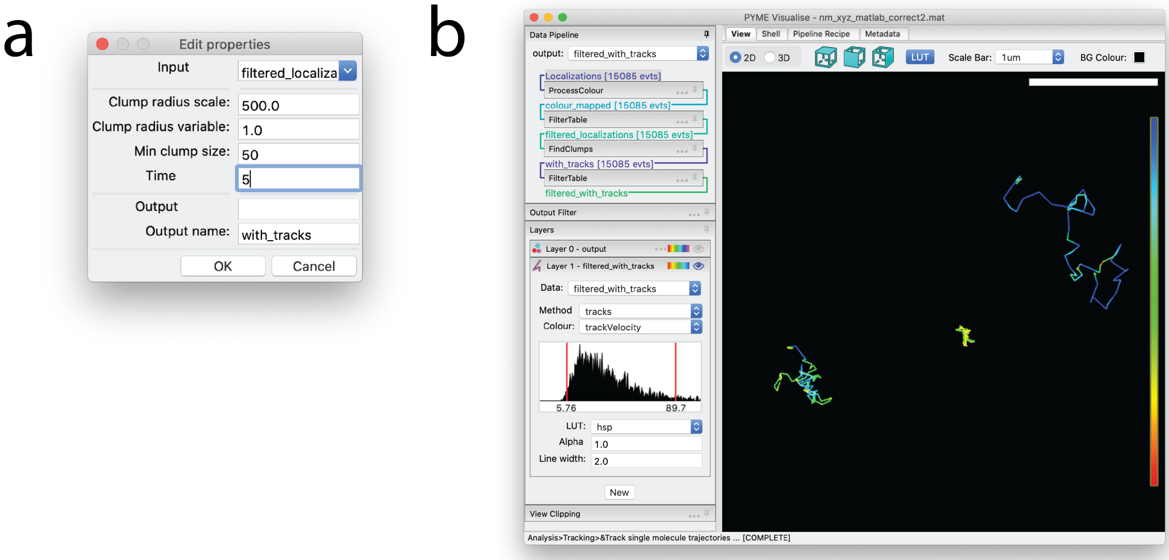

Fig. 6.2: Tracking Rtn4-SNAP in 2D. (a) Dialog window indicating settings for particle tracking. (b) The resulting trajectories, displayed with constant coloring. The particles giving rise to these trajectories are visible as points in Layer 0, which is hidden in this example, as indicated by the transparent eye.

#### 6.3 Other algorithms

**DBSCAN** An implementation of DBSCAN clustering ([ester1996], [nicovich2017]) is available by selecting *Analysis→Clustering→DBSCAN Clump* from the menu.

**QPAINT** Algorithms for ploting and fitting off-time distributions for QPAINT ([jungmann2016])

**Chromatic shift calibration** A couple of algorithms (*Corrections→Shiftmaps→XXX*) for calibrating chromatic shifts between colour channels from bead datasets. Mostly used with ratiometric localisation analysis.

**Vibration characterisation** Accessed as *Extras→Diagnostics→Plot vibration spectra*, this looks for signatures of instrument vibration in a (rapidly acquired) bead localisation series.

### 7 Animation

Animations, such as point cloud fly-throughs, are generated from keyframes set by the user in the animation panel. The animation panel is accessed by selecting *Extras→Animation* from the PYMEVisualize menus.

A populated *Animation* pane is shown in Fig. 7.1. Each row in the animation table represents a keyframe. A keyframe marks the end of a transition in the animation. A keyframe is set by rotating, translating, or zooming the data to have a desired look and then pressing *Add* in the *Animation* pane. Double-clicking on a row allows editing of the keyframe name, which can be used as a unique descriptor, and duration, which edits the length of the transition from the previous keyframe to this keyframe (note that the duration of the first keyframe is therefore a dummy variable). Selecting a row and pressing *Delete* removes the selected keyframe from the animation. Keyframes can be saved and loaded in a JSON format using *Save* and *Load*, respectively. All keyframes can be removed at once by pressing *Clear*.

When the *play* button is pressed, the animation will play in as a seamless transition through the keyframes in order. If *Capture* is pressed, a dialog will pop up and the user will choose a folder in which to save this animation as a series of image files. Expanding *Settings...* reveals a dropdown menu labeled *Export file type*, which allows the user to change the file type of the images exported to JPEG, PNG, or TIFF.

### 8 Synthetic data

For testing purposes, it can be helpful to generate synthetic data. PYMEVisualize lets users generate synthetic data using a simulated worm-like chain model (with the right settings this can produce reasonable analogues of a wide range of filamenteous structures from microtubules to tightly folded DNA), from an image, or from a text file containing a list of coordinates. First select *Extras→Synthetic Data→Configure* and choose the point generating source and set properties associated with the source, as shown in Fig. 8.1 a. Then select *Extras→Synthetic Data→Generate fluorophore positions and events*, and a simulated set of localization events is generated, as shown in Fig. 8.1 b. This data can then be analyzed in the same way that real data would be.

To simulate additional localizations from the same set of points, but using different simulation parameters, edit the properties as in Fig. 8.1 a and then select *Extras→Synthetic Data→Generate events*.

### 9 Editing the pipeline "recipe"

As mentioned in *The data pipeline*, data flows through a configurable pipeline. When constructing a complex workflow (e.g. processing different colour channels in different ways - see *Isolating a single channel for processing*) it can be useful to edit this pipeline directly using the recipe editor (Fig. 9.1), accessible via the *Pipeline Recipe* tab. This also lets you access additional functionality such as feature based tracking, which is not currently accessible through the menus. The pipeline is specified by a recipe (a .json textual description) which describes the various processing steps to be applied. The recipe can be modified, either graphically or as text, saved and reloaded on a new dataset, or applied to multiple input files in a batch using the `bakeshop` utility.

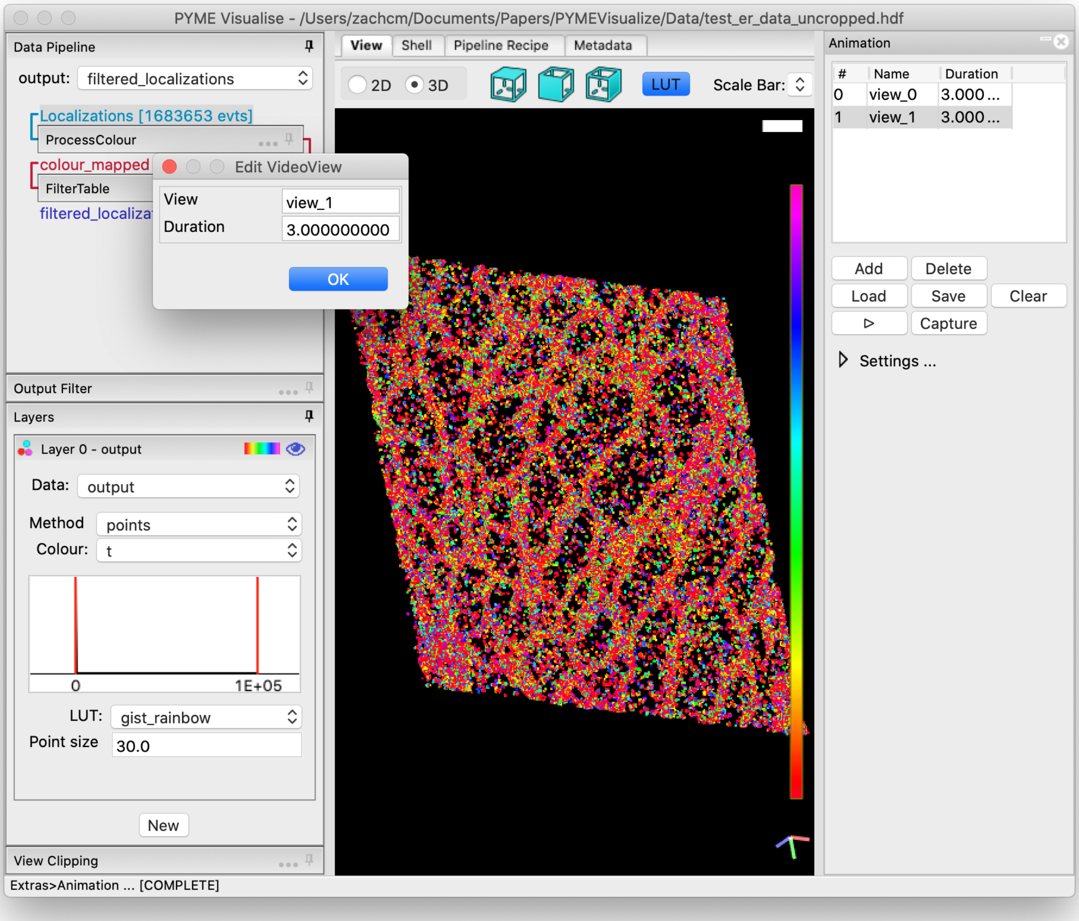

Fig. 7.1: Actively editing an animation keyframe in PYMEVisualize. This shows an animation with two keyframes added by rotating the data and pressing *Add* at each desired rotation. The second keyframe has been double-clicked, revealing an *Edit VideoView* window, which allows the user to change the name of the keyframe and the duration of the transition from the previous keyframe to this keyframe.

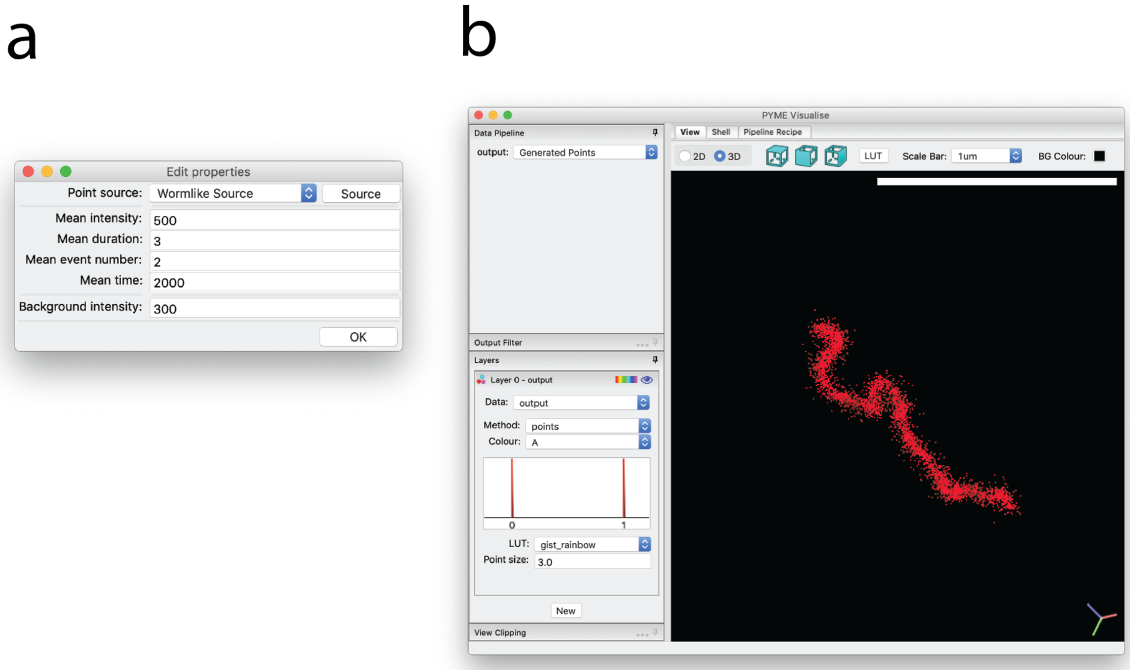

Fig. 8.1: Generation of synthetic data. (a) Dialog box that appears after selecting *Extras*→*Synthetic Data*→*Configure*. (b) Example synthetic worm-like chain created using parameters from dialog box in a.

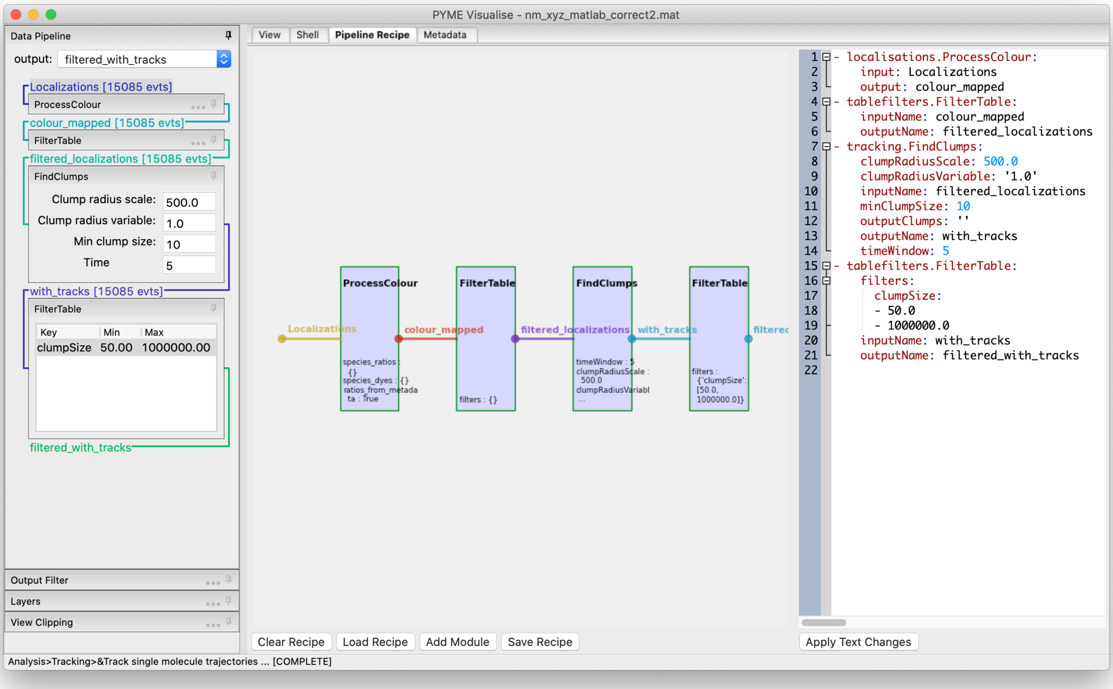

Fig. 9.1: The pipeline as seen in the recipe editor. Custom workflows can be created by manually adding processing modules.

### 10 Programmatic usage

#### 10.1 Shell

The *Shell* tab is a functional Python command line embedded within the program. The pipeline can be accessed directly from the shell, and behaves like a dictionary keyed by variable names. Pylab is imported in the shell making a number of MATLAB-style plotting and basic numeric commands accessible (see the matplotlib webpage for more docs). One can, for example, plot a histogram of point amplitudes by executing `hist(pipeline['A'])`. Pipeline data sources can be accessed by entering `pipeline.dataSources[datasource_key]`. For a list of datasource keys, type `pipeline.dataSources`.

#### 10.2 Jupyter notebook

PYMEVisualize can be used directly from a Jupyter notebook. At the top of the notebook, enter `from PYME.LMVis import VisGUI`, `%gui wx`, and then `pymevis = VisGUI.ipython_pymevisualize()`. This makes it possible to access a PYMEVisualize instance from a notebook through the `pymevis` variable. Setting `pipeline=pymevis.pipeline` gives the user access to the PYMEVisualize pipeline in exactly the same way as described in *Shell* section. An example of generating a point cloud, passing it to a tabular data source, and visualizing the data source is shown in Fig. 10.1.

```
In [ ]: from PYME.LMVis import VisGUI

        %gui wx
```

```
In [ ]: pymevis = VisGUI.ipython_pymevisualize()
        pipeline = pymevis.pipeline
```

```
In [ ]: import numpy as np
        from PYME.IO import tabular

        points = np.random.randn(100,3)*100
        pipeline.addDataSource('points', tabular.mappingFilter({'x': points[:,0],
                                                                'y': points[:,1],
                                                                'z': points[:,2]}))
        pipeline.selectDataSource('points')
        pymevis.add_pointcloud_layer(ds_name='points')
```

Fig. 10.1: An example of generating a point cloud, passing it to a tabular data source, and visualizing the data source in PYMEVisualize from a Jupyter notebook.

Please note that both PYME and the Jupyter kernel must be set up in the Framework build on a Mac (see https://python-microscopy.org/doc/Installation/InstallationFromSource.html) for this to work. To install the Jupyter kernel in the Framework build, activate `your_pyme_environment` and then type `PATH/TO/CONDA/ENVIRONMENT/python.app/Contents/MacOS/python -m ipykernel install --user --name your_pyme_environment`.

#### 10.3 Plugins

Details on extending and writing plugins for PYMEVisualize are available at http://python-microscopy.org/doc/hacking.html. A template for extending PYMEVisualize can be found at https://github.com/python-microscopy/pyme-plugin.

### 11 Further reading

Documentation is kept up-to-date at http://python-microscopy.org/doc/.

### 12 Appendix

#### 12.1 Ratiometric colour settings

When processing ratiometric localisation data (having a `gFrac` column) the splitting ratios for each dye species can either be set in the series metadata, or by using the *Colour* tab (Fig. 12.1). To add a labelling right click in the *Fluorophores* list box and select *Add*. Enter a channel name and splitting ratio in the dialog that opens. Alternatively click *Guess* to attempt to automatically detect the channels using the K-means algorithm. Once added, you can click in the name or ratio columns to edit. The plot above is a scatter plot showing a subset of all localisations and updates to show the resulting channel assignments.

Thresholds used in the Bayesian assignment process are adjustable in the *Channel Assignment* panel. A dye is assigned to a given channel if both it's probability of belonging to that channel is greater than `p_dye` **and** it's probability of belonging to any other channel is less than `p_other`. The defaults assign fluorophores to the most likely channel and ensure that the chance of a false assignment is less than 10%. We find they seldom need tweaking. If adjustment is necessary, `p_other`, which controls the rejection of potentially mis-assigned localisations, is most useful. It is tempting to think that `p_dye` should be higher (i.e. we should have a high certainty that a dye belongs to a given channel), but this would be a mistake - `p_dye = 0.1` will include 90% of the statistical spread of localisations belonging to that channel. `p_dye = 0.5` by comparison would only capture 50% of a dye's statistical spread and would needlessly discard a large fraction of the localisations.

#### 12.2 Isolating a single channel for processing

To apply processing steps to a single channel (rather than to all channels at once), it needs to be isolated in the pipeline. To do this, navigate to the *Pipeline Recipe* tab and select *Add Module*, as in Fig. 12.2 b. Then select the `ExtractTableChannel` recipe from the **localisations** recipes and press *Add*. This will result in a dialog box as shown in Fig. 12.2 c, where here the first color channel, `chan0`, is selected. Returning to the *View* tab and selecting `filtered` as the *output* in the upper-left portion of the window shows only the localizations present in the color channel `chan0` (Fig. 12.2 a, bottom). Additional data processing will only operate on this color channel as long as `filtered` is selected as the *output*.

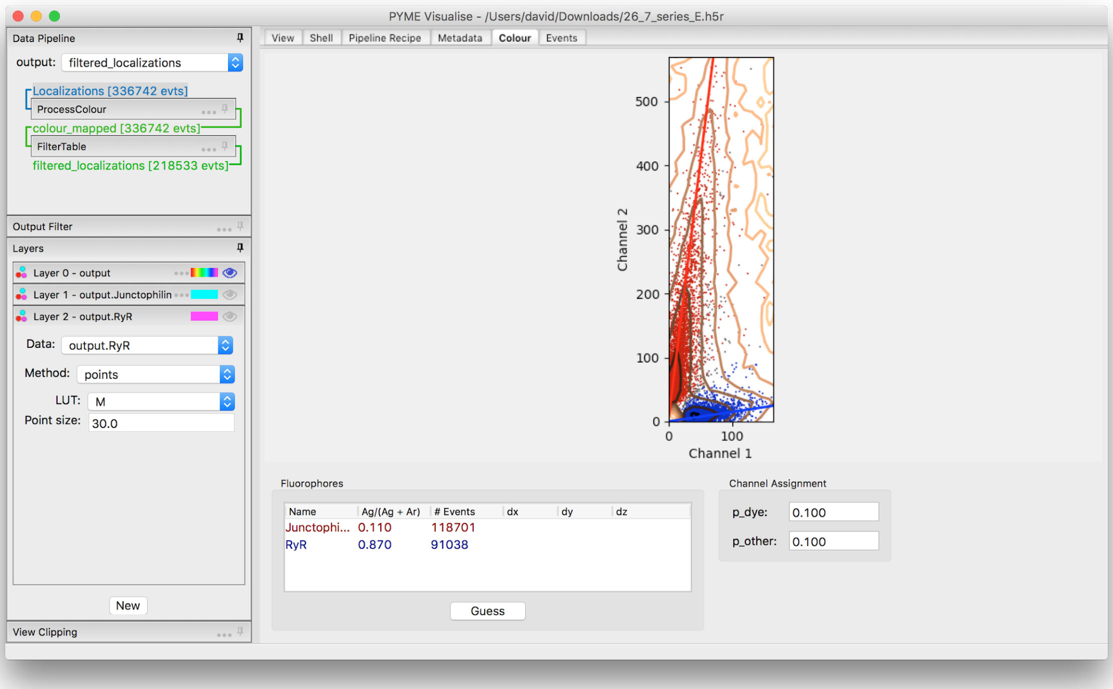

Fig. 12.1: Ratiometric splitting in the *Colour* tab.

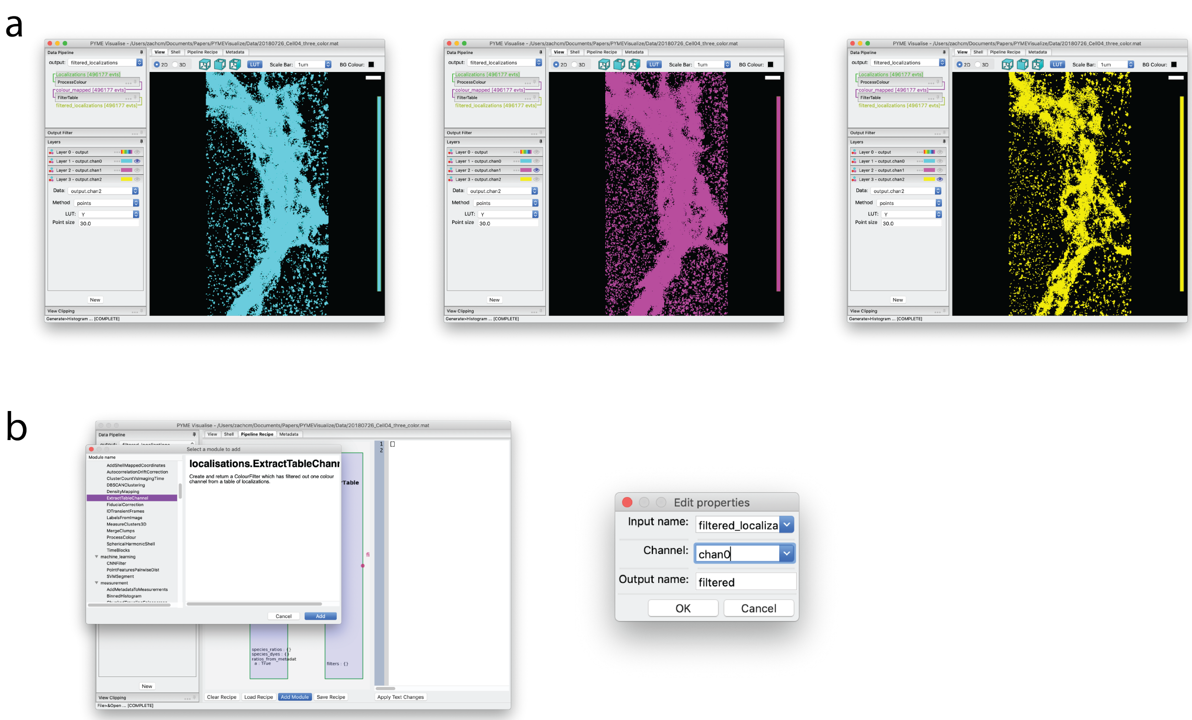

Fig. 12.2: Visualization and color channel selection of 3-color super-resolution image of *cis*, *medial*, and *trans*-Golgi. (a) *Top*. All three color channels visualized in a single layer. (b). Selection of the `ExtractTableChannel` recipe in the *Pipeline Recipe* tab and `ExtractTableChannel` dialog box, set to extract color channel `chan0` from the original data.
