## Supplementary note 2 for "PYMEVisualize: an open-source tool for exploring 3D super-resolution data"

### A quick tour of PYMEVisualize

In this tutorial, we'll touch on several aspects of the PYMEVisualize workflow, including opening a single-molecule localization microscopy (SMLM) data set, viewing it as points, filtering, and reconstructing density images, as well as extracting a 3D surface from the data set. The tutorial does not cover all (or even most) functionality, but rather aims to give a taste of what is possible. We assume PYMEVisualize is installed on your computer (see supplement or https://python-microscopy.org/doc/Installation/Installation.html), and that you have downloaded `test_er_data.hdf`, a supplementary SMLM data set of the endoplasmic reticulum provided with this paper.

#### Launch PYMEVisualize

For the purposes of this tutorial, we'll launch PYMEVisualize from a command line. On windows, open **Anaconda Prompt**. On Mac, open **Terminal**.[1] Once the command line is open, type `PYMEVis` and then press **enter** to launch the application[2].

Once launched, navigate to *File* > *Open*.

[1] If PYME is installed outside of the Anaconda base environment, type `conda activate <pyme_envronment>`. If you are unsure of where PYME is installed, assume you do not need to type this command.

[2] On Windows you should also be able to select **PYMEVisualise** from the start menu

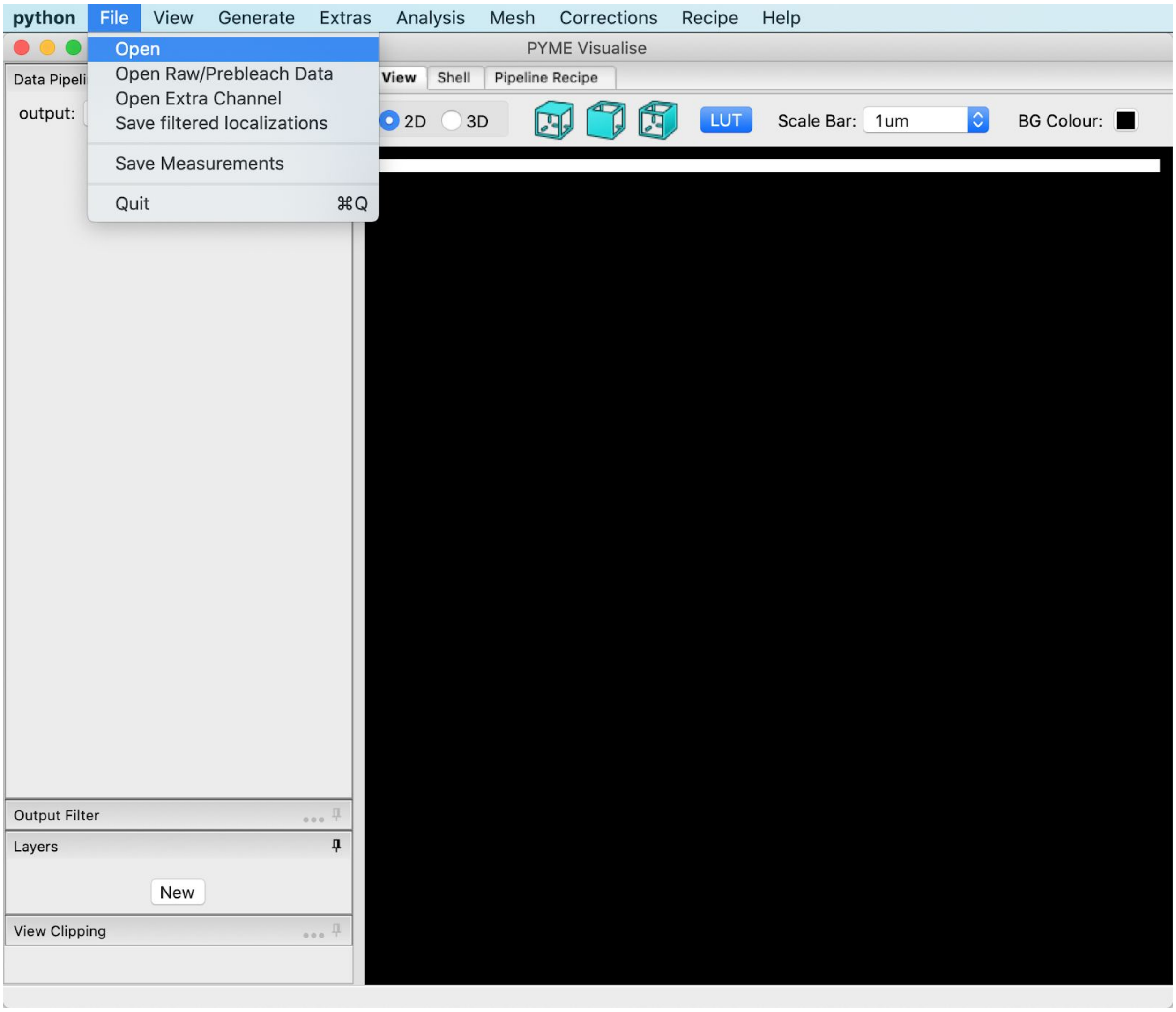


You will be prompted with a file dialog asking you to *Choose a file to open*. Select `test_er_data.hdf`, which is provided with this publication. Your screen will appear as below.

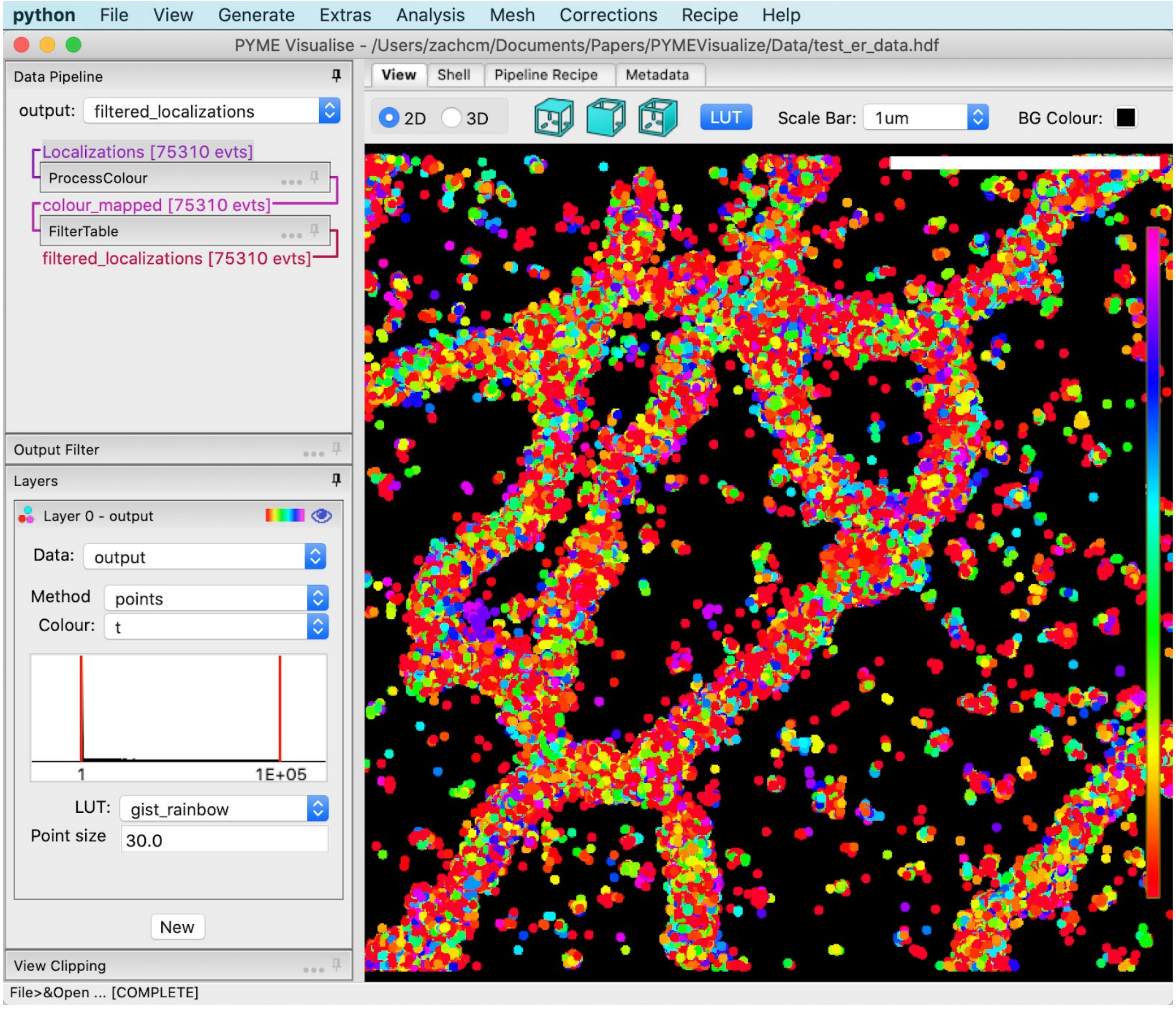


#### Filtering

We want to restrict our data to well localized events. Click the *FilterTable* box in the data pipeline view to expand, as shown below.

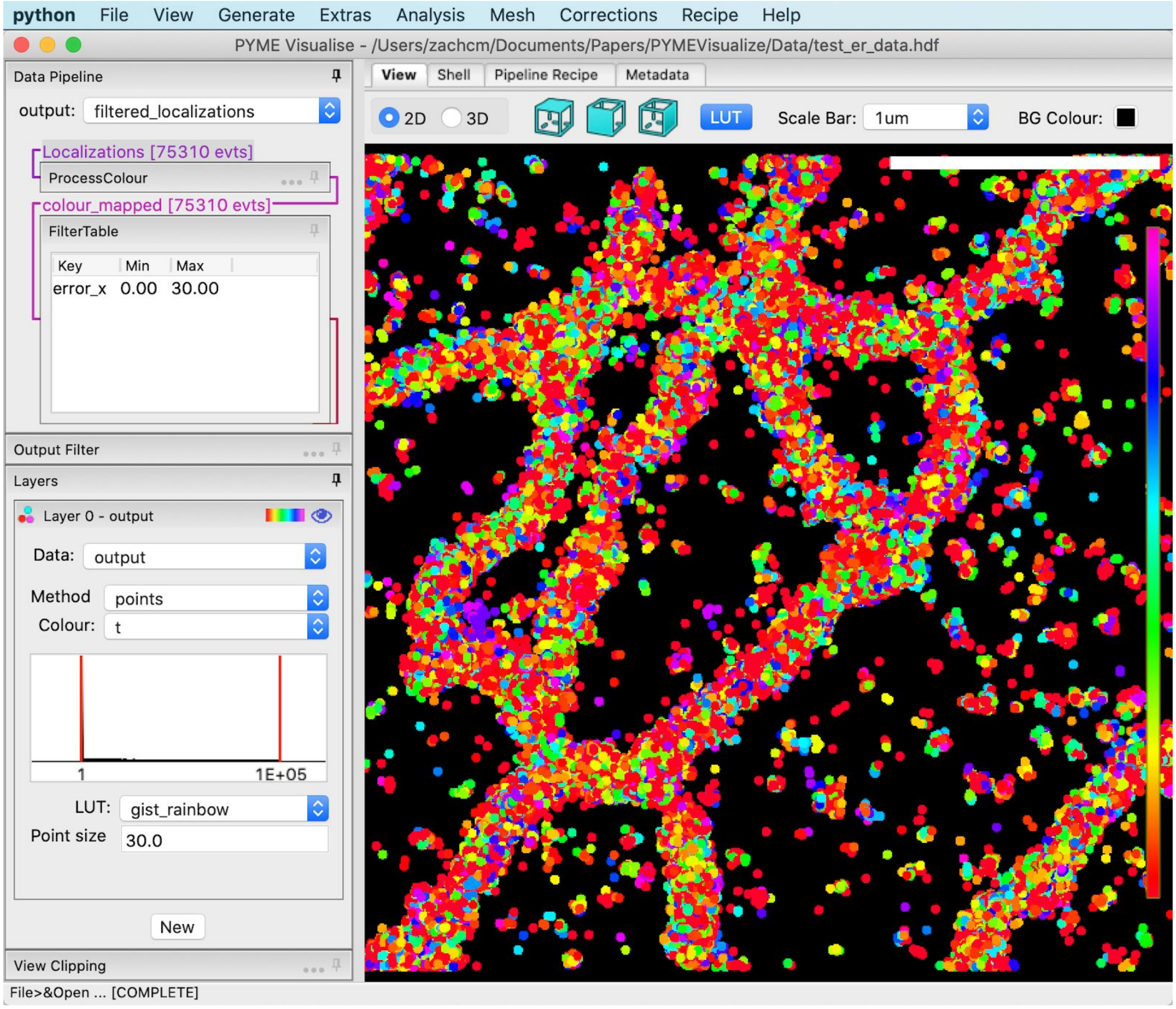


Double click on the entry for `error_x` to bring up the dialog as shown below. Drag the right hand red bar in the histogram towards the left to restrict the localisation error to the range from 0 to 10 nm. Press *OK*.

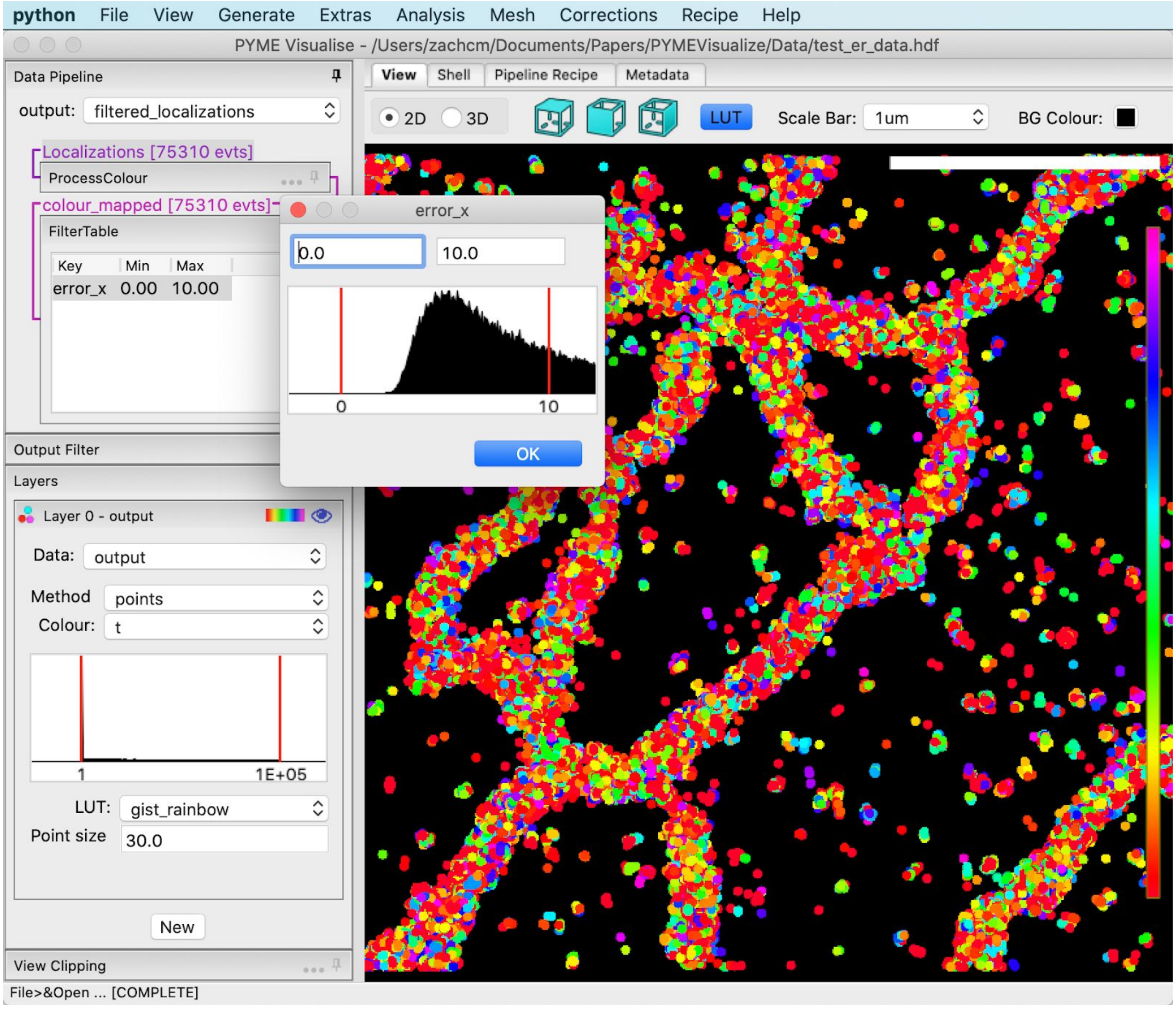


#### Adjust visualization parameters

By default, data opens with localisations coloured by time as this gives a quick visual indication if there is a problem with drift. The example dataset has negligible drift, so lets try some other options. Under *Layer 0 - output* Change the “Colour” to `z`, which is the axial position of the localisation, and pull the red bars on the histogram display under “Colour” in to adjust the colour scaling to exclude the outliers and make the depth changes in the structure more visible, as shown below. Alternatively, click in the histogram box and press **p** to ask PYMEVisualize to automatically set the histogram lower and upper bounds to 1st and 99th percentile of values, respectively .

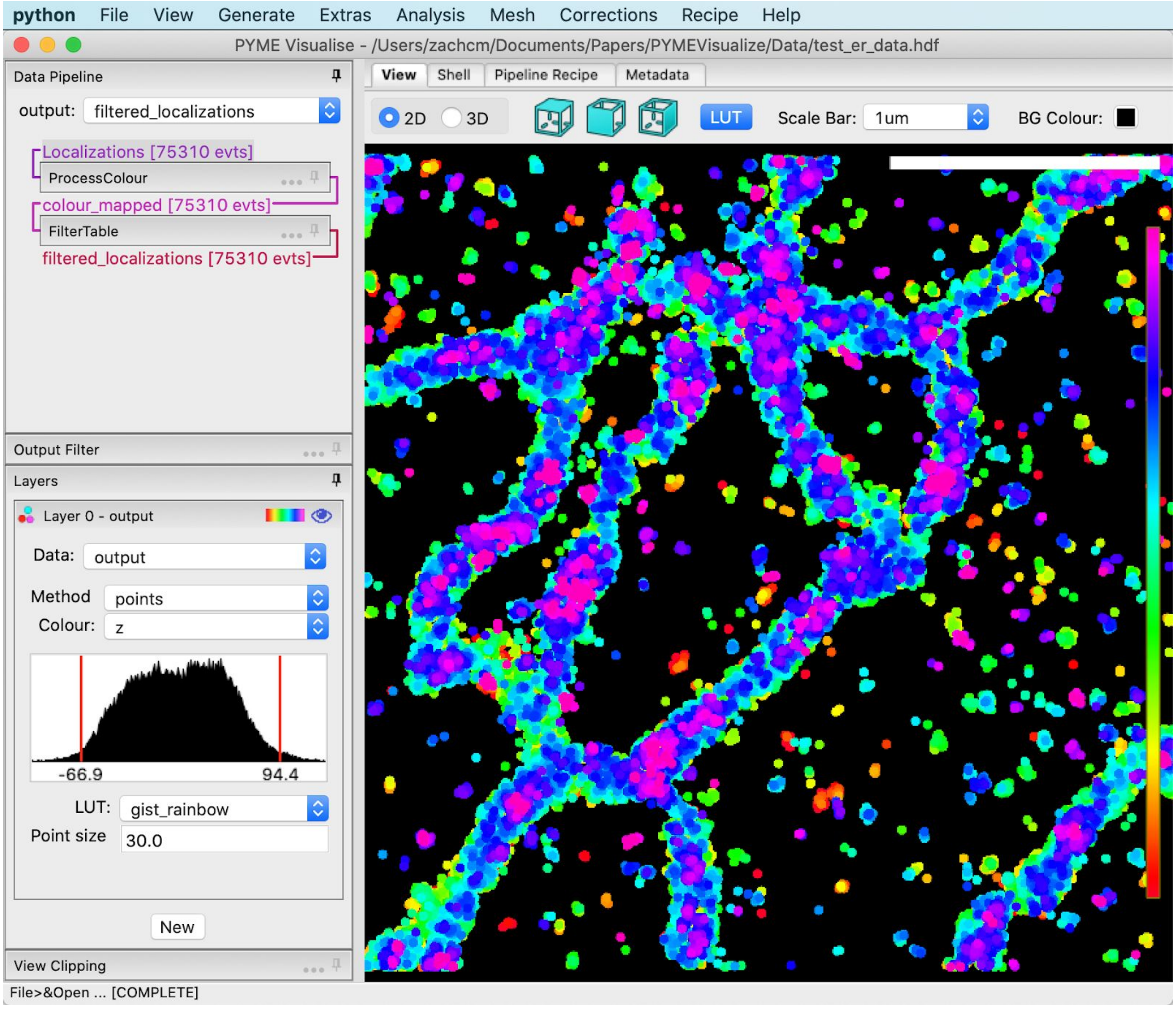


The simple `points` mode is very busy and tends to get swamped in areas of high point density. Switch “Method” to `pointsprites`, “Point size” to `10.0`, and “Alpha” to `0.2` to get a real-time approximation of the popular Gaussian reconstruction method. Switch “LUT” to `hsp` to get constant-intensity coloring.

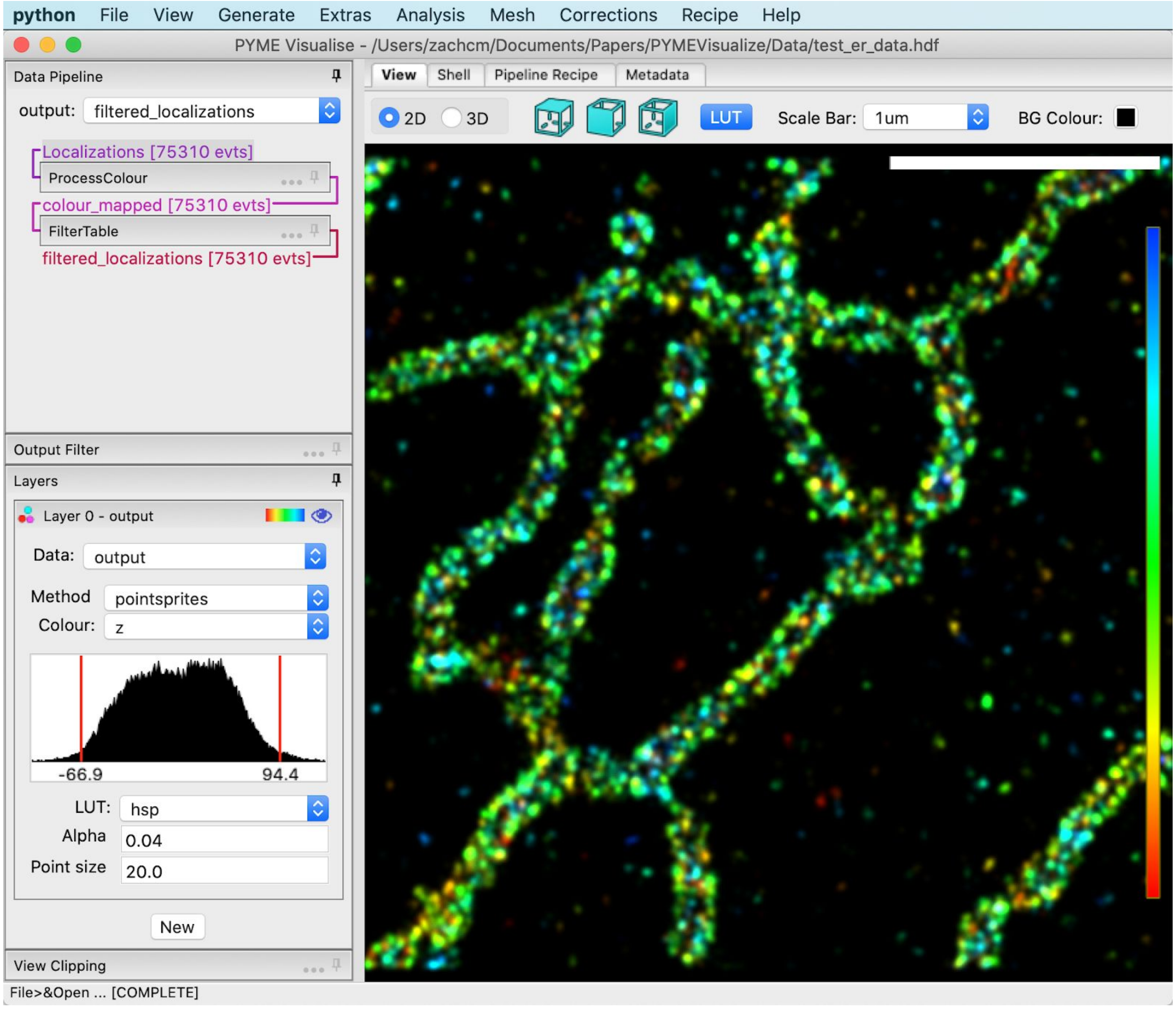


#### Generate a reconstruction

Let’s create a 2D histogram reconstruction of our data set that we could use for pixel-based analysis. Navigate to *Generate* > *Histogram*. A dialog box will pop up as shown below.

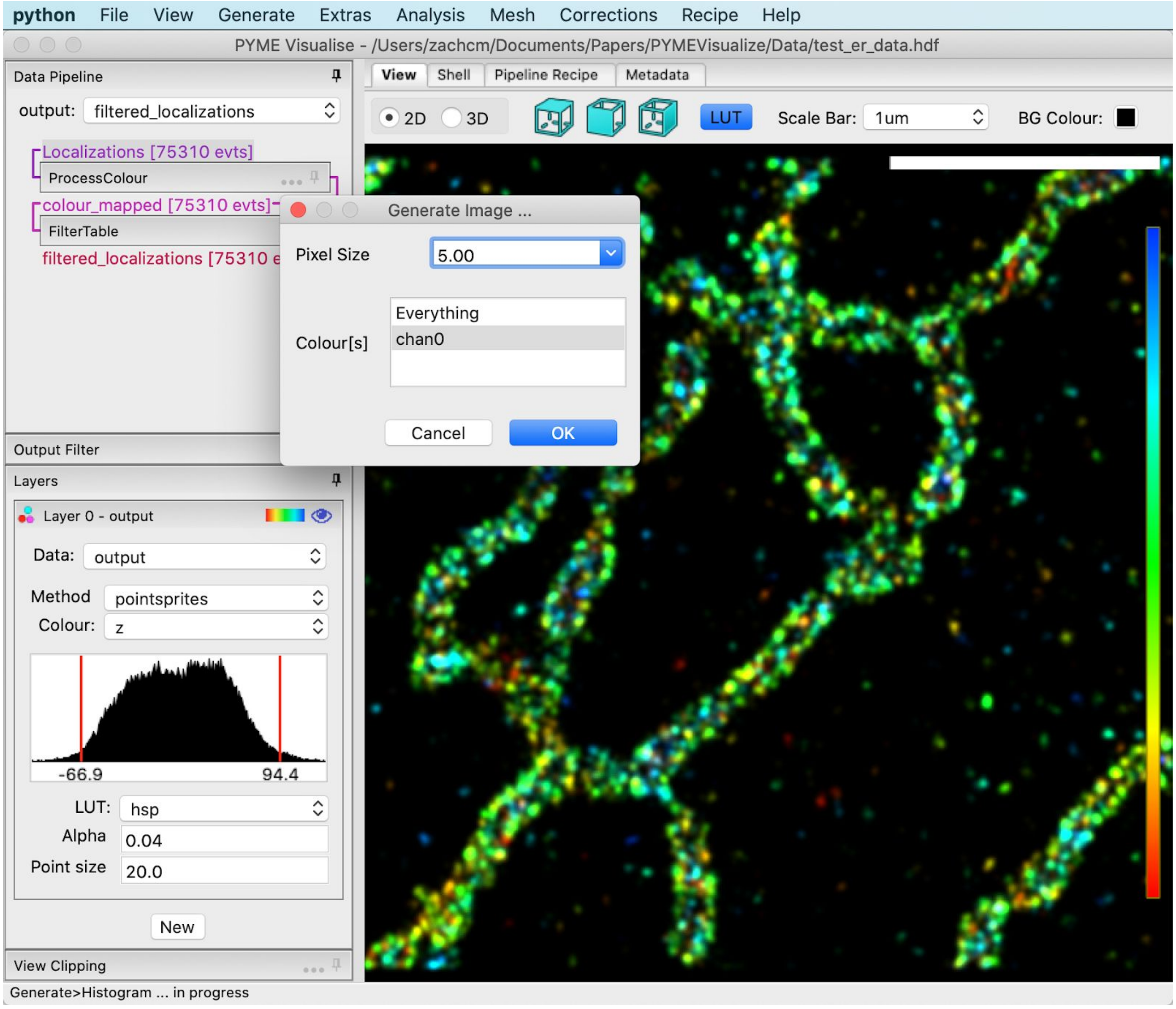


Notice that in the lower left corner of the window, it says “Generate > Histogram … in progress”. This area of the program lets the user know what is currently running and if it is completed.

Leave the “Generate Image …” dialog options as default and press *OK*. A 2D histogram image will appear, as shown below.

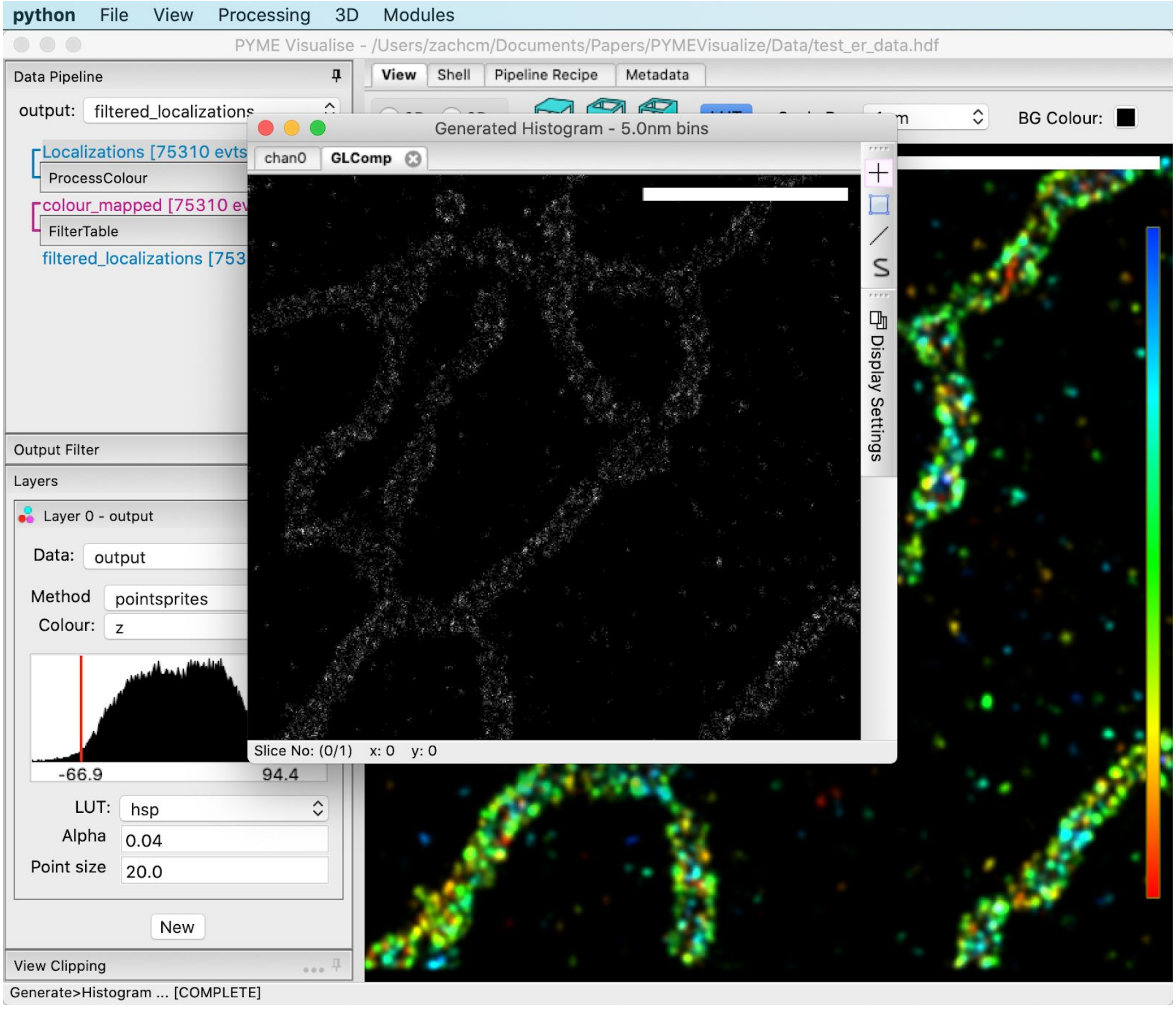


To adjust the contrast of the histogram displayed in the “GLComp” tab, click *Display Settings* on the right of the histogram window. Pull the red bars on the histogram display under “chan0” to adjust contrast.

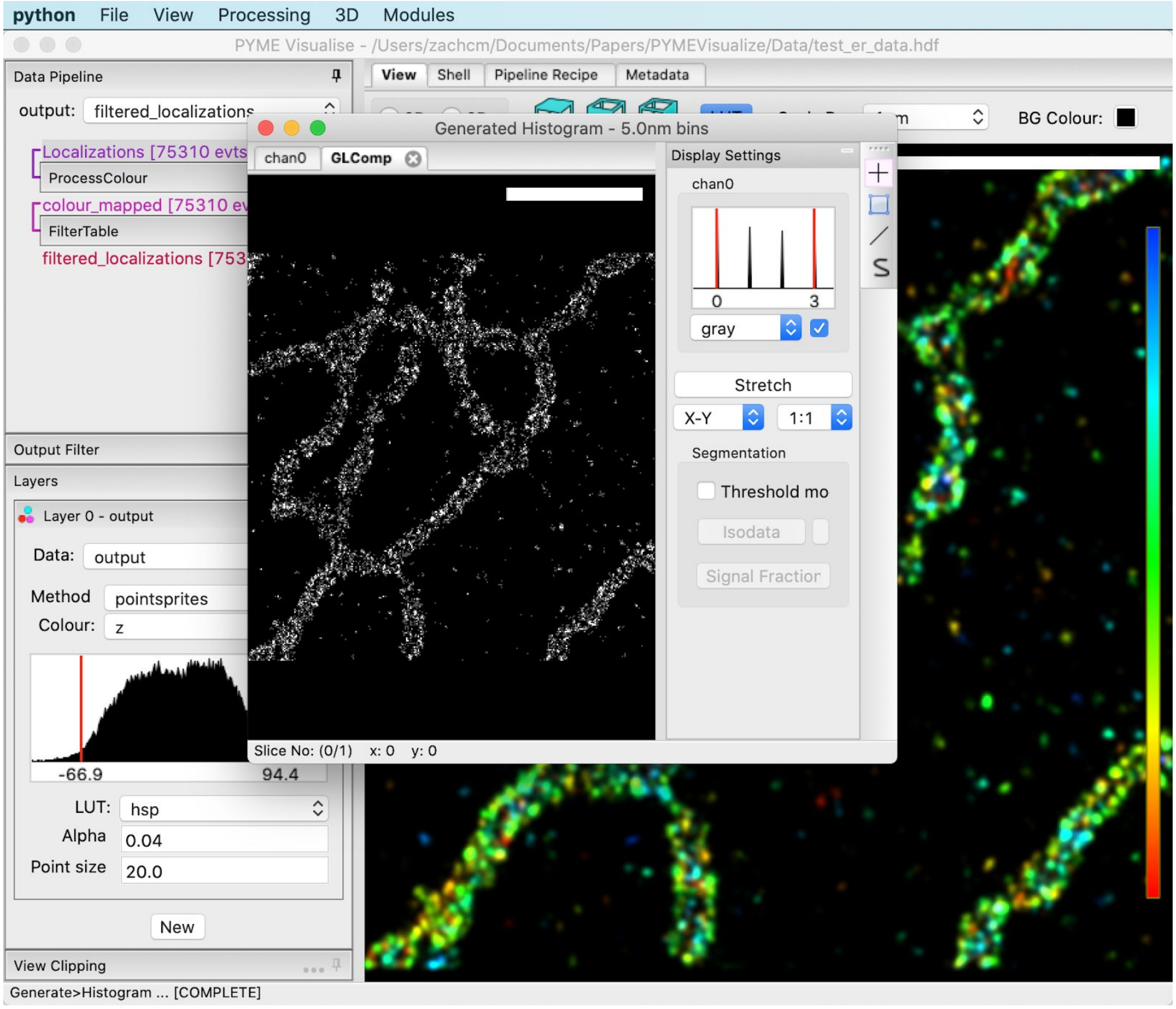


Navigate to *File* > *Save As* and name your file “histogram_rendering.tif”. This is a voxel-based image which can be opened and analysed using the same tools (e.g. ImageJ) that you might use for conventional microscopy images. The *Generate* menu also has options for a bunch of other 2D and 3D density reconstruction methods (see User Guide for details). Open in the image editor of your choice[3].

#### Explore data in 3D

Return to the main PYMEVisualize window, optionally closing the Gaussian rendering window. Select the “3D” radio button under the “View” tab near the top of the screen. Click on the data and drag with your mouse to rotate the data view. Right-click on the data and drag to translate

[3] Some versions of ImageJ/FIJI do not load floating point TIFF (and therefore our exported images) correctly, although the Bio-Formats importer does. If an exported .tif looks weird in ImageJ, try opening with the Bio-Formats importer.

the data view. Rotate the scroll wheel to zoom in and out of parts of the data. Choose a rotation, translation, and zoom that you think looks nice. Ours is below.

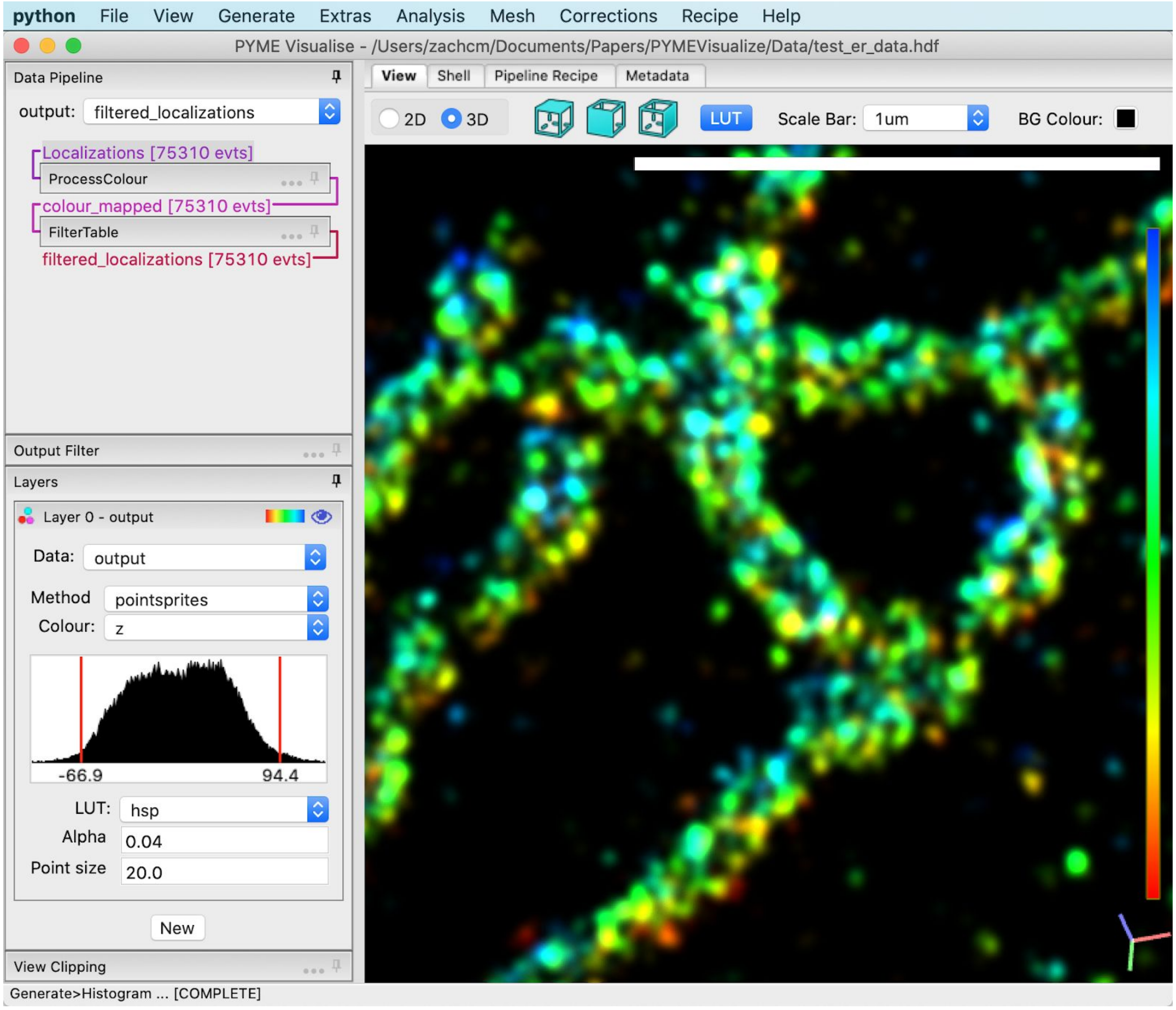


Since we've zoomed in, the 1 micrometer scale bar is looking rather big. Change the scale bar size to "200nm", as shown below.

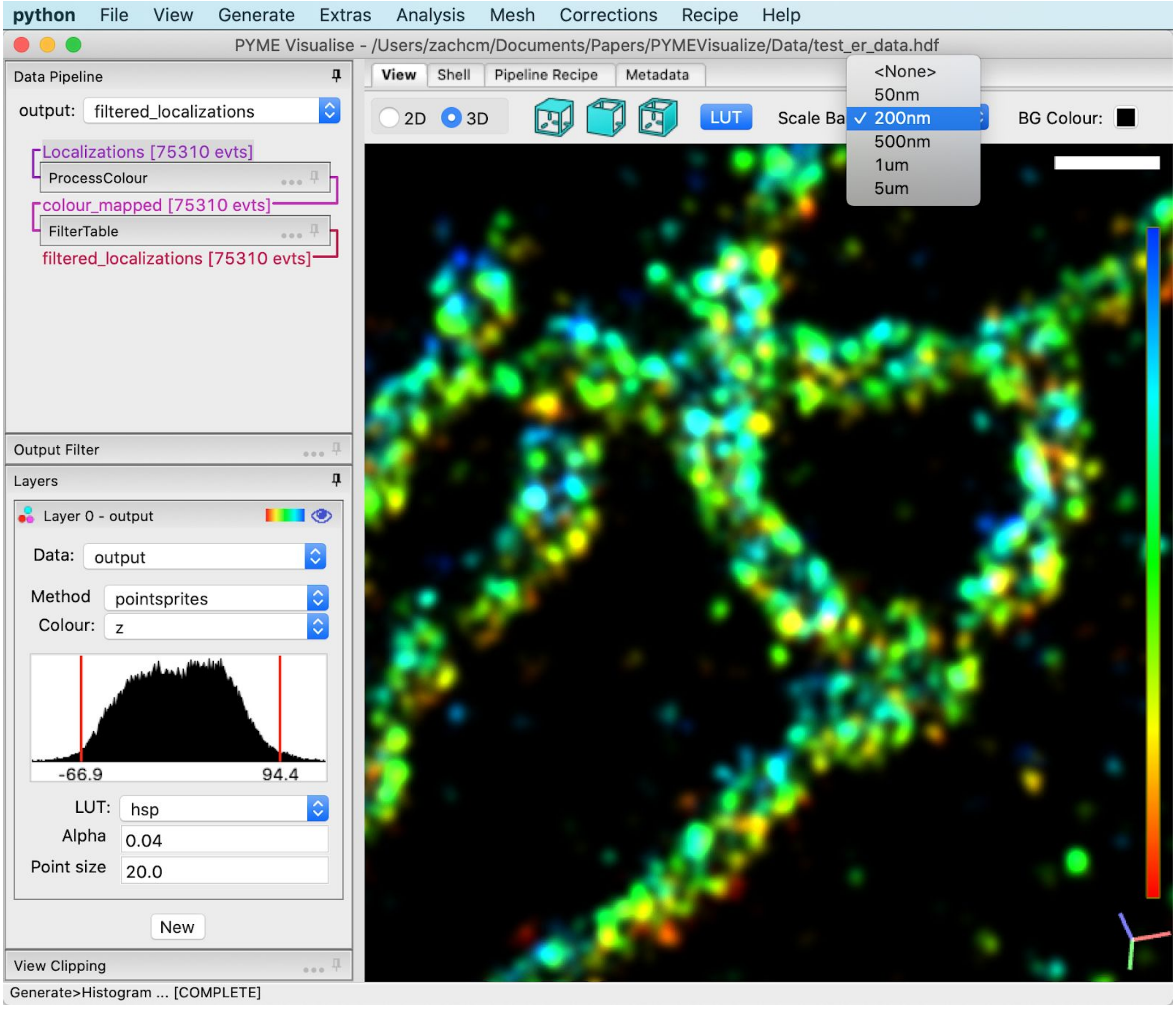


#### Generate an isosurface from point cloud data

This is a data set of membrane-bound proteins on the endoplasmic reticulum. As such, it is a good approximation of the membrane surface. We can generate an approximation of the ER's surface from this data by navigating to *Mesh > Generate Isosurface*. A dialog box will appear. Change the properties to what is shown in the image below, then press *OK*.

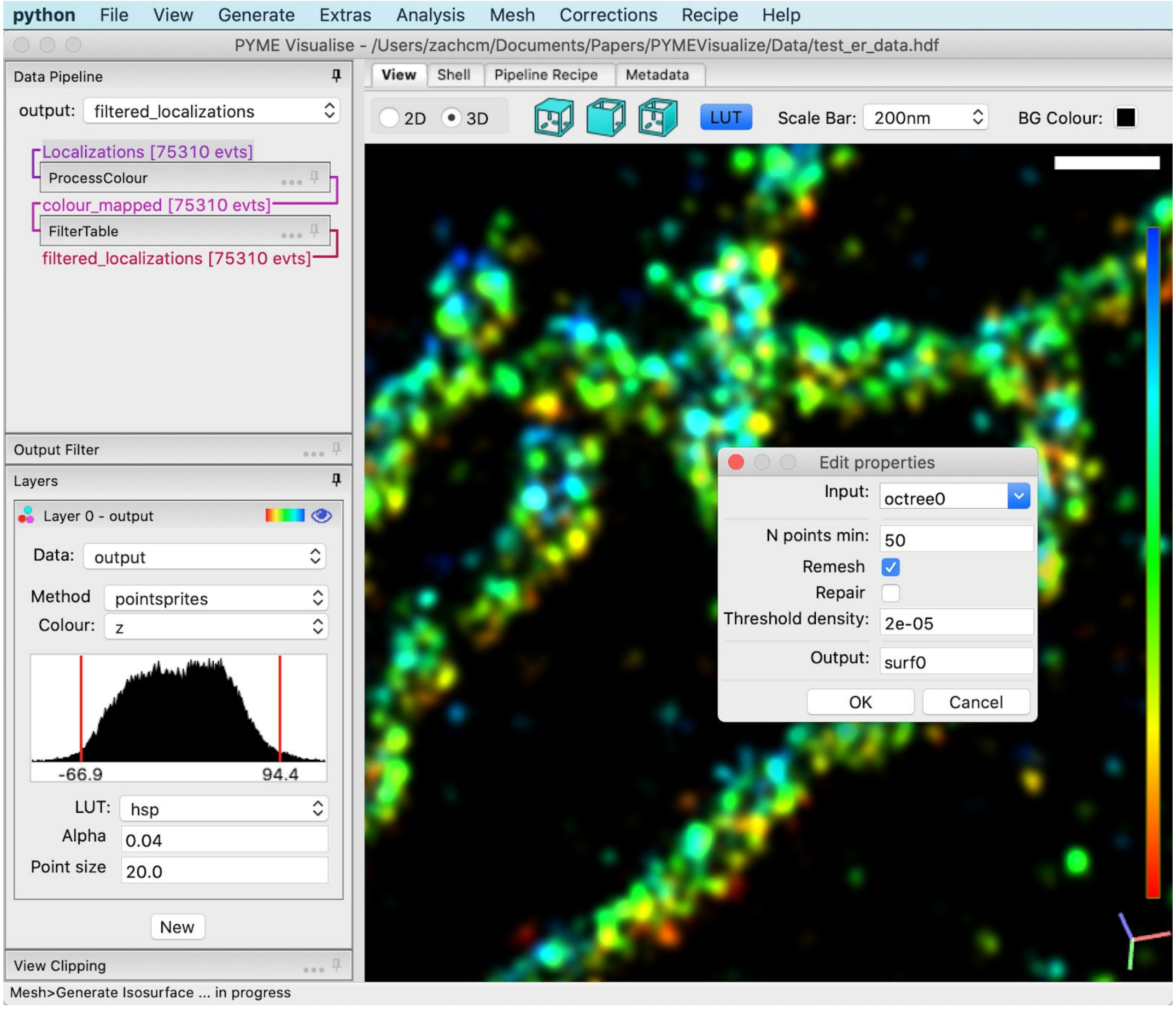


Give this 30 seconds to run. Once you see “Mesh>Generate Isosurface … [COMPLETE]” in the lower left corner, you are ready to proceed to the next step. The window should appear as below.

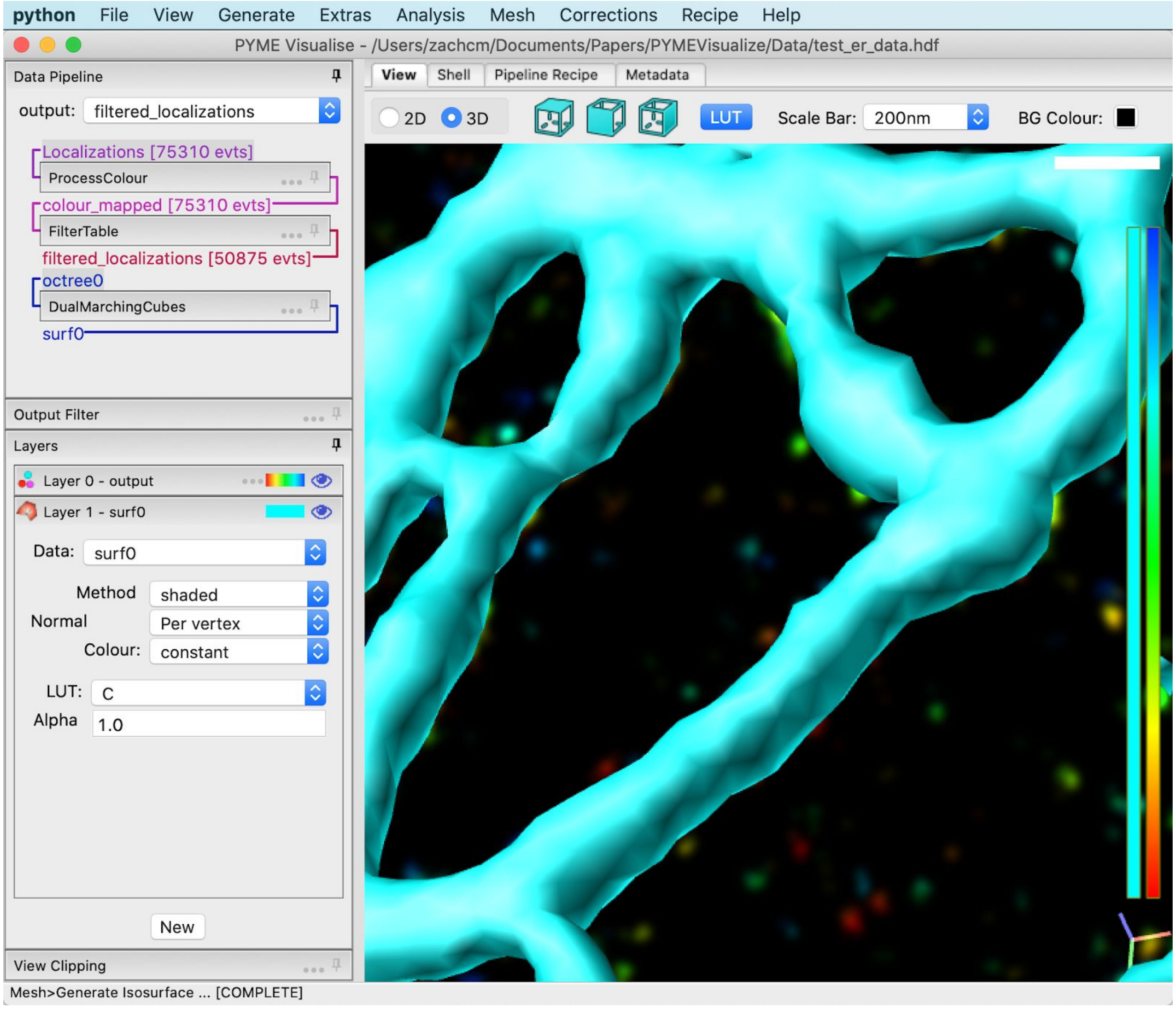


Notice that we now have two “Layers”. The first is *Layer 0 - output*. We played with this layer’s parameters in the “Adjust visualization parameters” section earlier. *Layer 0 - output* is called a points layer, as indicated by its three colorful points. The second layer is called *Layer 1 - surf0*, and it is for displaying surfaces, as indicated by its red triangles. Layers stack on top of one another in the viewing area. As with the point layer, you can change how the surface layers appear - try changing “Alpha” in *Layer 1 - surf0* to `0.5`. You should see something similar to the image below.

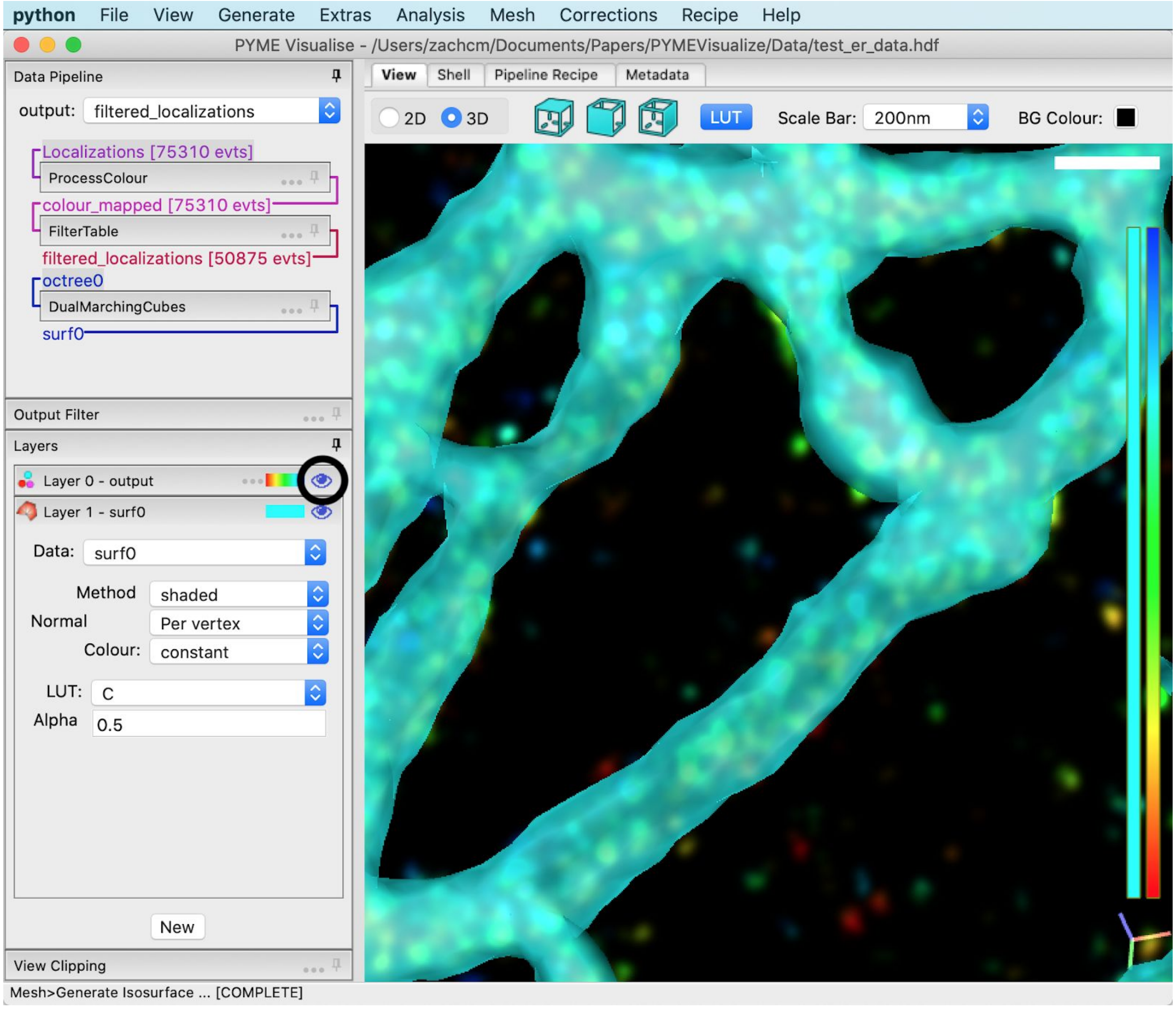


The visibility of individual layers can be toggled using the eye button associated with it. Press on the eye associated with *Layer 0 - output*, circled in black above.

#### Color the isosurface by mean curvature

Like points, surfaces can be coloured by a number of different parameters. Let's color this isosurface by its mean curvature. Surfaces start off using solid colour lookup tables (one of 'C', 'Y', 'M, 'R, 'G', 'B') which display the same colour regardless of what the colour variable is, so the first thing we need to do is change the "LUT" to something which will show differences in our colour value. "RdBu" is a good choice in this case. Now, change "Colour" to "curvature_mean", as shown below.

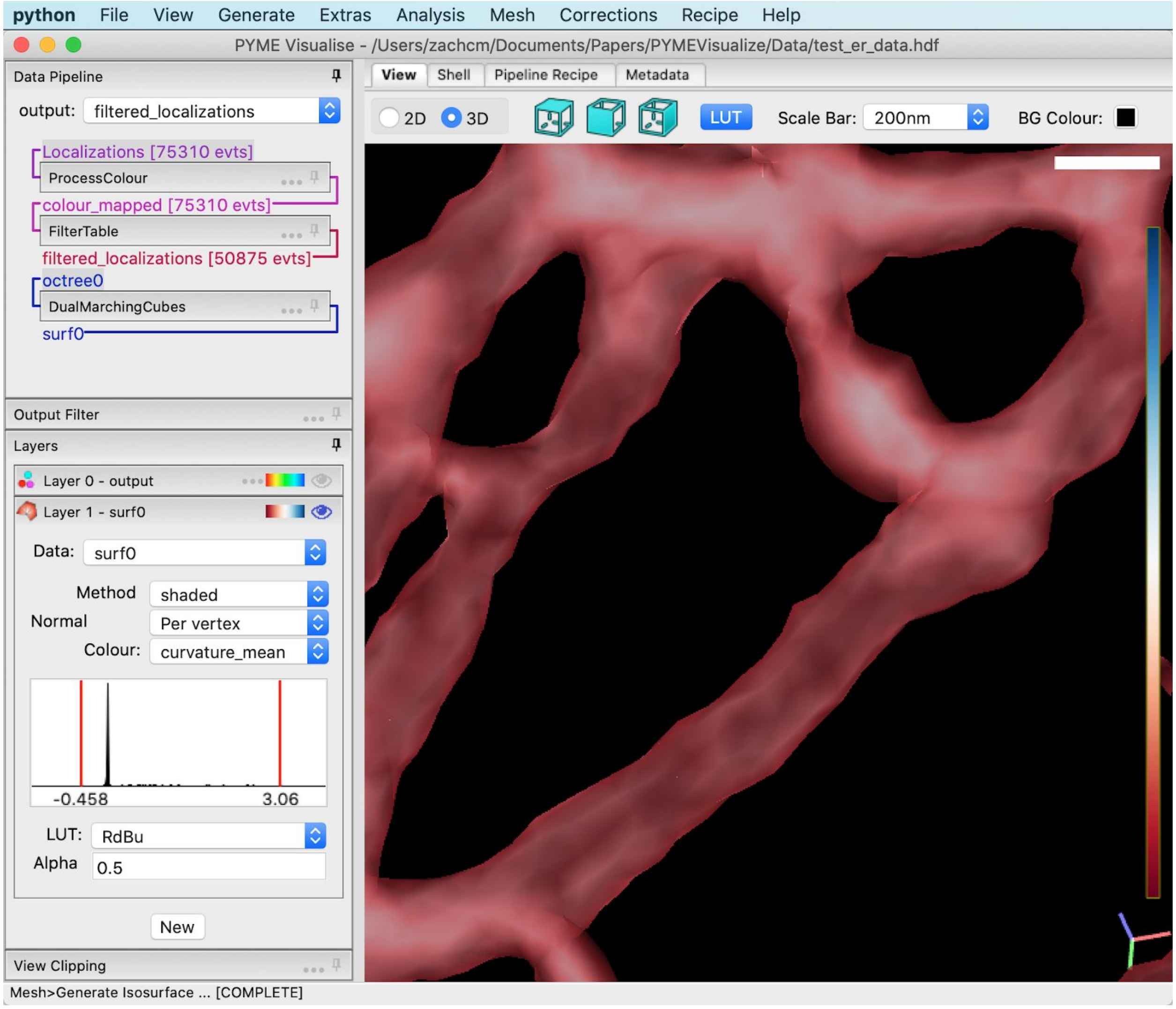


We don't see a lot of colour in the result, as we have a few outliers in our curvature estimates which broaden the distribution of values so that all the interesting curvature values map to one LUT point. We can change this using the histogram below "Colour". Right click in the middle of it to get a dialog box, and change the dialog box parameters to read `-0.02` for "Min" and `0.02` for "Max". Our mean curvature is expressed in units of 1/nanometer, making this range correspond to realistic physiological curvatures ≤ 1/50nm.

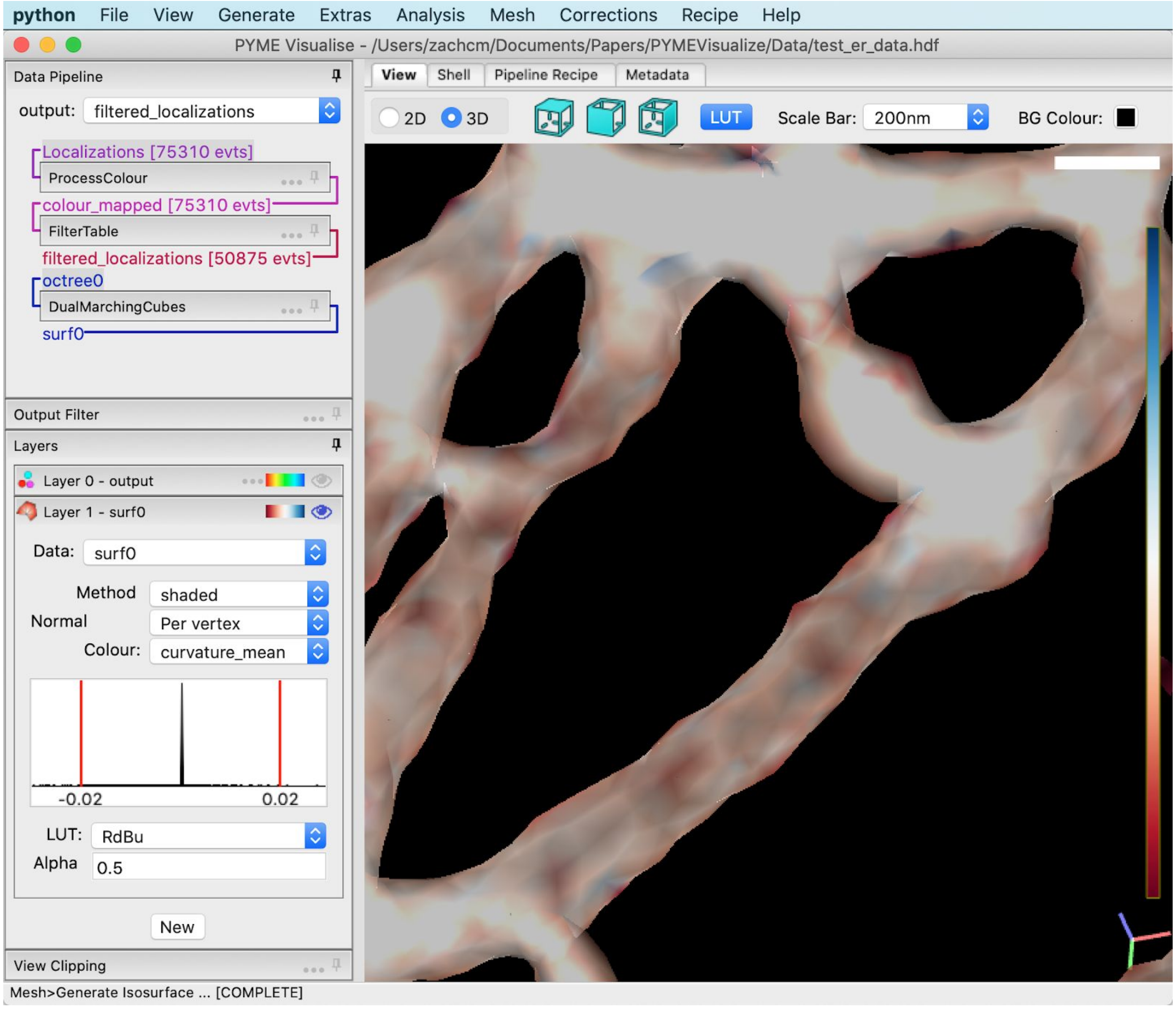


The surface is now colored by mean curvatures between -0.02 $nm^{-1}$ and 0.02 $nm^{-1}$. As expected, flat regions of the surface are white (close to 0 nm), curved surfaces are red (closer to 1/50 nm), and dips in the surface are blue (closer to -1/50 nm).

As before, we can rotate and zoom the view. Also try toggling the LUT using the toolbar button. Once you have a view you are happy with, export a snapshot using the *View >Save snapshot* menu item. This should give you a .png which can be viewed with standard image viewers or embedded in Word, PowerPoint, Illustrator, and other publication and presentation tools.

#### Conclusion

You can now open, visualize, and export 2D and 3D data, but this barely scratches the surface of PYMEVisualize's capabilities. To discover (or develop) additional features, see the User Guide or visit python-microscopy.org.
